# CeLINC, a fluorescence-based protein-protein interaction assay in *C. elegans*

**DOI:** 10.1101/2021.06.01.446599

**Authors:** Jason R Kroll, Sanne Remmelzwaal, Mike Boxem

## Abstract

Interactions among proteins are fundamental for life and determining whether two particular proteins physically interact can be essential for fully understanding a protein’s function. We present *C. elegans* light-induced co-clustering (CeLINC), an optical binary protein-protein interaction assay to determine whether two proteins interact *in vivo*. Based on CRY2/CIB1 light-dependent oligomerization, CeLINC can rapidly and unambiguously identify protein-protein interactions between pairs of fluorescently tagged proteins. A fluorescently tagged bait protein is captured using a nanobody directed against the fluorescent protein (GFP or mCherry) and brought into artificial clusters within the cell. Co-localization of a fluorescently tagged prey protein in the cluster indicates a protein interaction. We tested the system with an array of positive and negative reference protein pairs. Assay performance was extremely robust with no false positives detected in the negative reference pairs. We then used the system to test for interactions among apical and basolateral polarity regulators. We confirmed interactions seen between PAR-6, PKC-3, and PAR-3, but observed no physical interactions among the basolateral Scribble module proteins LET-413, DLG-1, and LGL-1. We have generated a plasmid toolkit that allows use of custom promoters or CRY2 variants to promote flexibility of the system. The CeLINC assay is a powerful and rapid technique that can be widely applied in *C. elegans* due to the universal plasmids that can be used with existing fluorescently tagged strains without need for additional cloning or genetic modification of the genome.

**Summary:** We have developed a protein-protein interaction assay for *C. elegans* to investigate whether pairs of proteins interact *in vivo*. *C. elegans* light-induced co-clustering (CeLINC) is based on trapping a fluorescently-tagged bait protein into artificial clusters, and observing whether candidate interacting prey proteins co-cluster with the bait protein. CeLINC can be widely applied as a single set of universal plasmids can be used with existing strains expressing fluorescently-tagged proteins.

## Introduction

Interactions among proteins are critical for the functioning of the cell. Characterizing protein-protein interactions (PPIs) is therefore necessary to understand protein function, and numerous technologies have been developed to identify PPIs. One commonly used technique is the yeast two-hybrid system (Walhout et al., 2000), which allows for high-throughput screening, but takes place in a context different from the original organism or cell type. Affinity purification combined with mass spectrometry can identify multiple targets interacting with a protein of interest (Dunham et al., 2012) but tissue-specific information is lost and animals are analyzed in bulk. PPI assays that can be applied *in vivo* often rely on split or complementary tags that assemble upon physical proximity of the two proteins to be tested (Bischof et al., 2018; Brückner et al., 2009; Hudry et al., 2011; Shyu et al., 2008), and generally require the generation of fusion proteins that have no uses outside of the interaction assay. To complement these existing assays, we have adapted a fluorescence-based PPI assay for use in *C. elegans*, CeLINC, that can utilize existing fluorescently tagged strains, is easily analyzed, and can be performed in single animals in any cell type of interest without using specialized equipment.

CeLINC is based on a method originally developed in mammalian cell culture (Taslimi et al., 2014) and is an extension of an optogenetic protein inhibition system called “light-activated reversible inhibition by assembled trap” (LARIAT). LARIAT can inhibit target proteins in living cells in a spatiotemporally controlled manner by sequestering the target protein into clusters (Lee et al., 2014). LARIAT elegantly exploits the cryptochrome 2 (CRY2) protein that homodimerizes and heterodimerizes with the cryptochrome-interacting basic-helix-loop-helix (CIB1) protein upon blue light exposure (Kennedy et al., 2010). By fusing a target protein to CRY2 or through the use of nanobodies, target proteins can be sequestered and inhibited in a light-dependent manner. Together, this system has been used to inhibit proteins in a variety of pathways (Asakawa et al., 2020; Nguyen et al., 2016; Qin et al., 2017; Qin et al., 2018) and to recently control mRNA localization (Kim et al., 2020).

With the LARIAT system as the basis, light-induced co-clustering (LINC) has been developed as a binary PPI assay (Osswald et al., 2019; Taslimi et al., 2014; Ventura et al., 2020). In LINC, two proteins tagged with different fluorescent proteins are tested for their ability to co-cluster in the blue-light-induced CRY2/CIB1 clusters (Figure 1A). To modularize the assay, a GFP nanobody is attached to CRY2 to allow recruitment of any GFP-tagged bait protein to CRY2 clusters in blue light (Figure 1A). After cluster formation, co-clustering of a prey protein tagged with a differently colored fluorescent protein (*e.g.,* mCherry, mScarlet, BFP, or mKate2) is assessed. Proteins that show a positive protein interaction show co-localization in the clusters, while proteins that do not interact do not co-localize in the clusters (Figure 1B). LINC is analogous to a typical immunoprecipitation experiment but takes place within the living cell and allows for visual identification of the protein interaction. The assay therefore requires minimal equipment, only relies on fluorescent protein tags, can maintain cell type specificity, and can be easily scored without complex analysis. Additionally, the use of a nanobody gives the system flexibility since any fluorescently tagged protein with a suitable nanobody epitope can be used as the bait protein without additional modifications.

**Figure 1.**
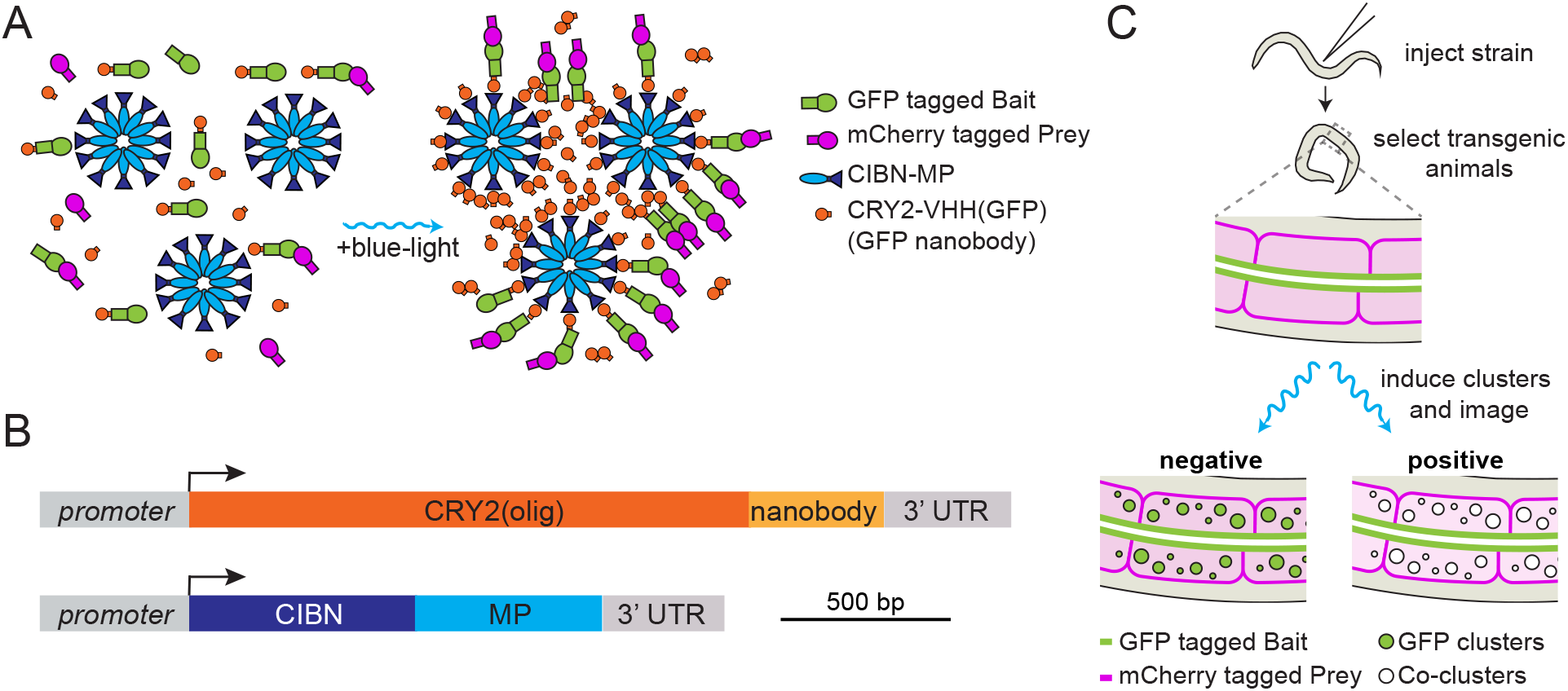
Overview of CeLINC. (**A**) Overview of CRY2/CIB1 light-induced co-clustering (LINC). Two proteins to be tested for interaction are tagged with fluorescent proteins (GFP and a second color fluorescent protein). In dark conditions, the CRY2::VHH(GFP nanobody) protein is mainly in the non-oligomerized form, and there is no to little association between CIBN with CRY2. Upon blue light exposure, CRY2 both homodimerizes and heterodimerizes with CIBN. The CIBN-MP protein forms a dodecamer that act as a scaffold to increase cluster size. GFP-tagged proteins bound to the nanobody are clustered, resulting in a bright and compact fluorescent signal. The second color fluorescent protein (mCherry in this example) is analyzed for co-localization in the clusters. (**B**) Overview of CeLINC. A strain with two tagged proteins to be tested for an interaction (GFP and mCherry in this example) is injected with plasmids to express the CRY2::VHH(GFP nanobody) and CIBN-MP proteins. Transgenic F1 animals carrying an extrachromosomal array are identified by the presence of a co-injection marker. After transgenic strains are established, clusters are induced by blue light exposure of the transgenic animals and imaged. Cells with GFP containing clusters are then analyzed for co-localization of the mCherry tagged protein. (**C**) Diagram of the CeLINC expression constructs.

CRY2 based oligomerization and clustering has been used previously in *C. elegans* to induce aggregation of Amyloid-β to study how aggregates affect Alzheimer’s disease pathologies, and to oligomerize UNC-40 to manipulate growth cone development, but not to investigate PPIs (Endo et al., 2016; Lim et al., 2020). We adapted LINC for *C. elegans* (CeLINC) and have validated its function and utility to investigate PPIs. First, we have adapted components of the system by codon-optimizing the proteins for use in *C. elegans* and have created expression plasmids that allow for expression of CRY2/CIB1 proteins in multiple tissues. Next, we have characterized CRY2/CIB1 clustering characteristics upon blue light exposure in various *C. elegans* tissues. We then tested CeLINC on various positive and negative reference protein pairs. Finally, we tested for interactions among cell polarity regulators due to their extensively studied nature and previously established protein interactions in other systems and *C. elegans*. We provide a plasmid toolkit to enable flexibility and adaptability of the CeLINC system for further studies and uses. Due to the universal nature of the plasmids and the ability to use existing fluorescent strains, the CeLINC system is an extremely rapid and powerful way to characterize PPIs in *C. elegans*.

## Results

### Light-induced CRY2 clustering in *C. elegans*

We designed the CeLINC system as a two-vector system (Figure 1B). One vector expresses a fusion of a nanobody (VHH) with a variant of the CRY2 protein, CRY2(olig) (E490G), that increases oligomerization (Taslimi et al., 2014). The other vector expresses the CIB1 N-terminal region (CIBN) fused to the multimerization domain (MP) of Ca2+/Calmodulin-dependent protein kinase II (CaMKII). Inclusion of MP enhances light-activated oligomerization and helps to increase cluster sizes by acting as a scaffold (Che et al., 2015; Lee et al., 2014). All of the components were codon optimized for expression in *C. elegans*. We first designed vectors for expression with a general promotor, *rps-0*, which expresses broadly in somatic cells including the intestine, muscle cells, and hypodermis. Constructs were assembled modularly with the SapTrap assembly system (Schwartz and Jorgensen, 2016) to allow for further flexibility and ease of use for future modifications or variants. To use CeLINC, in brief, animals expressing fluorescently tagged proteins to be tested for interaction are injected with the CeLINC plasmids to form extrachromosomal arrays (Figure 1C). Transgenic strains are established by selecting animals expressing a co-injection marker, and strains are established that reliably transmit the extrachromosomal array. Finally, animals are exposed to blue light and are imaged to determine whether the two fluorescently tagged proteins co-cluster in the cell types of interest or not, indicating a positive or negative protein interaction respectively (Figure 1C).

We first tested the ability for CRY2(olig) to form clusters in response to blue light. To directly visualize cluster formation, we tagged CRY2(olig) with the fluorescent protein mKate2. Worms were injected with two plasmids: mKate2::CRY2(olig)::VHH(GFP), and a protein fusion of CIBN::MP, both expressed from the *rps-0* general promoter and using the *unc-54* 3’ UTR. We used a non-fluorescent co-injection marker that confers a Roller (Rol) phenotype and resistance to hygromycin to identify animals carrying extrachromosomal arrays. Transgenic strains were kept in complete darkness during development, mounted on slides under dark lighting conditions, and then placed on the spinning disk microscope in near darkness, with the dimmest amount of light possible to center the worm on the microscope and focus (dark lighting condition). Before blue light stimulation, few to no mKate2::CRY2(olig) clusters were apparent in the animals, but a diffuse mKate2 signal was detected, corresponding to mKate2::CRY2(olig) proteins in a non-oligomerized state (Figure 2A). Next, blue light was delivered in a series of pulses to the animal (see Methods). Immediately after this treatment, mKate2::CRY2(olig) was found to be highly clustered in the cell, indicating that the CRY2 clustering system responds to blue light activation and readily and rapidly forms clusters, as seen in other systems (Figure 2A). Both intestinal and muscle cells showed rapid cluster formation (Figure 2A), in addition to other cells in the animal. Clusters were visible and had formed in different compartments of the cell, such as the nucleus, cytoplasm, and alongside the plasma membrane (Figure 2A). Overall, cellular morphology in the animals appeared normal, and there were no apparent phenotypes observed in the transgenic progeny. Therefore, blue-light-activated CRY2 clusters form rapidly and readily in *C. elegans*, and there was little to no toxicity or lethality associated with expression of the constructs.

**Figure 2.**
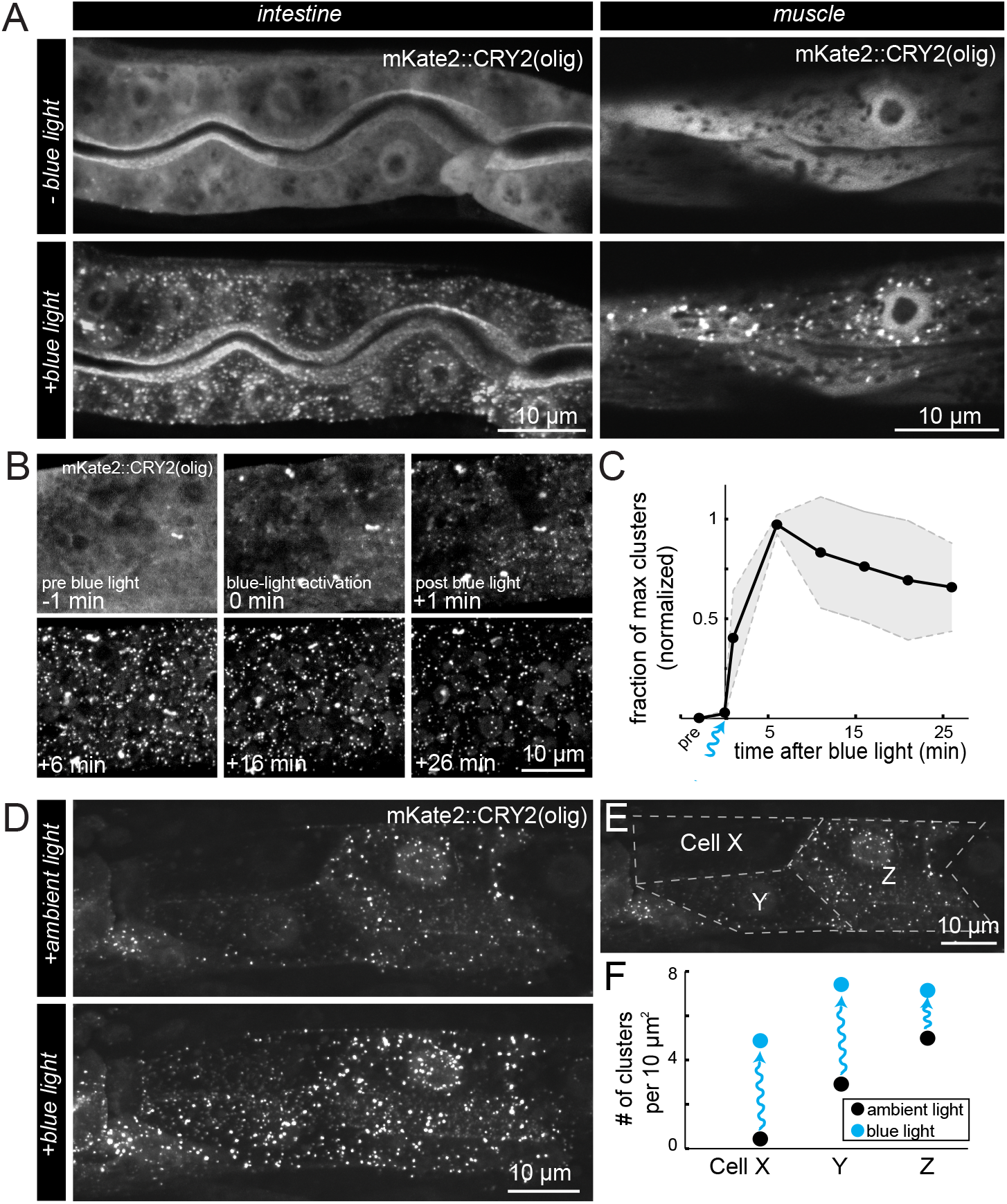
Behavior of CRY2(olig) clusters and light activation in *C. elegans.* (**A**) Fluorescent images (maximum z-stack projections) of intestine and muscle cells in transgenic animals expressing mKate2::CRY2(olig) and CIBN-MP before and after blue light exposure. mKate2 signal was diffuse before blue light exposure, while extensive clusters rapidly formed throughout the cytoplasm and in the nuclei upon exposure. Time between conditions was 1 minute. (**B**) Maximum projections of a region of an intestinal cell taken before (−1 min), during (0 min), and after (+1 min to +26 min) blue light activation to quantify oligomerization and decay of mKate2::CRY2(olig) clusters. (**C**) Quantification of cluster number over time in intestinal cells before and after blue light exposure, N=3 animals. Clusters were quantified with ComDet plugin for ImageJ/FIJI (see Methods). Black line indicates the mean fraction of clusters at a given time point, normalized to 0 before illumination and 1 at the point of maximum cluster formation. Grey shaded regions indicate the standard deviation. (**D**) Maximum projection images of the anterior portion of the intestine after ambient light exposure during mounting (normal binocular microscope with white light), and after additional blue light stimulation. (**E**) Schematic of the manual segmentation of the intestinal cells of the animal in (D) for quantification in (F). (**F**) The number of clusters per 10 μm^2^ in each cell segmented in (E), in ambient light and blue light conditions. Clusters were quantified with ComDet plugin.

To determine the half-life of the CRY2(olig) clusters in *C. elegans*, we grew and mounted animals in dark conditions, stimulated cluster formation with blue light, then imaged the animals over time with no further blue light stimulation (Figure 1B). We found that the maximum number of clusters was obtained six minutes after blue light exposure, and the decay in the number of clusters was slow, giving an estimated half-life of 34 min (Figure 1C). The rate of decay was roughly comparable to experiments in cell culture using a CRY2(olig)-mCherry construct, which showed a half-life of 23 min (Taslimi et al., 2014). These values are significantly longer than wild-type CRY2, for which a half-life of around 6 minutes has been reported (Bugaj et al., 2013; Lee et al., 2014). Therefore, CRY2(olig) clusters in *C. elegans* are relatively stable and allow ample time for animal manipulation and imaging.

Since it is inconvenient to manipulate animals in complete darkness and under special lighting conditions, in subsequent experiments we mounted animals on slides under ambient room lighting conditions and with white light from a binocular microscope. We expected this approach to pre-activate cluster formation before imaging. Pre-activation of clusters during mounting should also aid imaging, since animals and cells expressing the CeLINC constructs in the cell types of interest can be more quickly identified. In this condition, cells already displayed clustering of mKate2::CRY2(olig) before blue light stimulation, indicating that the white light received during mounting was sufficient to activate CRY2 oligomerization (Figure 1D). These animals were then subjected to blue light stimulation to determine whether additional oligomerization could be induced. Indeed, cells showed increased mKate2::CRY2(olig) clustering in response to blue light stimulation (Figure 1D). One such animal showed differing levels of mKate2::CRY2(olig) expression in three different intestinal cells, as determined by their level of diffuse mKate2 signal in the nucleus (Figure 1E, F). The cell with the weakest level of mKate2::CRY2(olig) expression showed no cluster formation under ambient light, but significant cluster formation after blue light stimulation, while the cells with higher amounts of mKate2::CRY2(olig) expression had less of a change after blue light exposure (Figure 1F). In addition, the area and number of clusters correlated to the mKate2::CRY2(olig) expression level (Figure 1F). Therefore, as seen in another study, higher expression levels induce clustering more readily than lower expression levels (Che et al., 2015). In summary, the blue-light-activated CRY2 based oligomerization behaves similarly to other systems, suggesting that the system is functional and suitable for use in *C. elegans*.

### Use of CeLINC for identifying PPIs

Next, after establishing CRY2 functionality and cluster formation in *C. elegans*, we tested the ability of the CeLINC assay to discriminate between positive and negative protein-protein interactions. We used the highly conserved and well-studied protein interaction between PKC-3 (aPKC) and PAR-6 (Par6). Both proteins are essential for polarization of cells in various tissues and form a complex through interaction of their PB1 domains (Li et al., 2010a). Mutations that delete this domain are lethal, highlighting the critical importance of this protein interaction during development and cell function (Li et al., 2010a).

To test this protein pair for physical interaction, we used a strain with an endogenous CRISPR/ Cas9 tagged *GFP::pkc-3* allele, and provided either *mKate2::par-6* or *mKate2::par-6*(*ΔPB1*), containing a small deletion of the PB1 domain (amino acids 15–28), to test for positive and negative interactions, respectively (Figure 3A, B). Both *par-6* variants were expressed with a general promoter from an extrachromosomal array to circumvent the lethality of mutated *par-6* alleles. Unlabeled *CRY2(olig)::VHH(GFP)* and the *CIBN-MP* plasmids were injected alongside either *mKate2::par-6* or *mKate2::par-6*(*ΔPB1)*. Clusters were pre-activated with white light during mounting, and we analyzed both the epidermal and the intestinal tissues. We found the wild-type protein pair showed extensive co-localization of GFP and mKate2 signal in cytoplasmic clusters (Figure 3C), while there was clearly a lack of co-localization in the PAR-6 mutant protein pair, in which the mKate2 signal remained diffuse (Figure 3D). Additionally, GFP::PKC-3 containing clusters showed on average a more than two-fold higher signal than the endogenous fluorescent signal of GFP::PKC-3 at the apical intestine (Figure 3E, F). The concentration and increased signal of the fluorescent protein in the clusters is a significant benefit for the CeLINC system, since it increases the visibility and signal of weakly expressed proteins, allowing them to be more easily identified for co-localization analysis.

**Figure 3.**
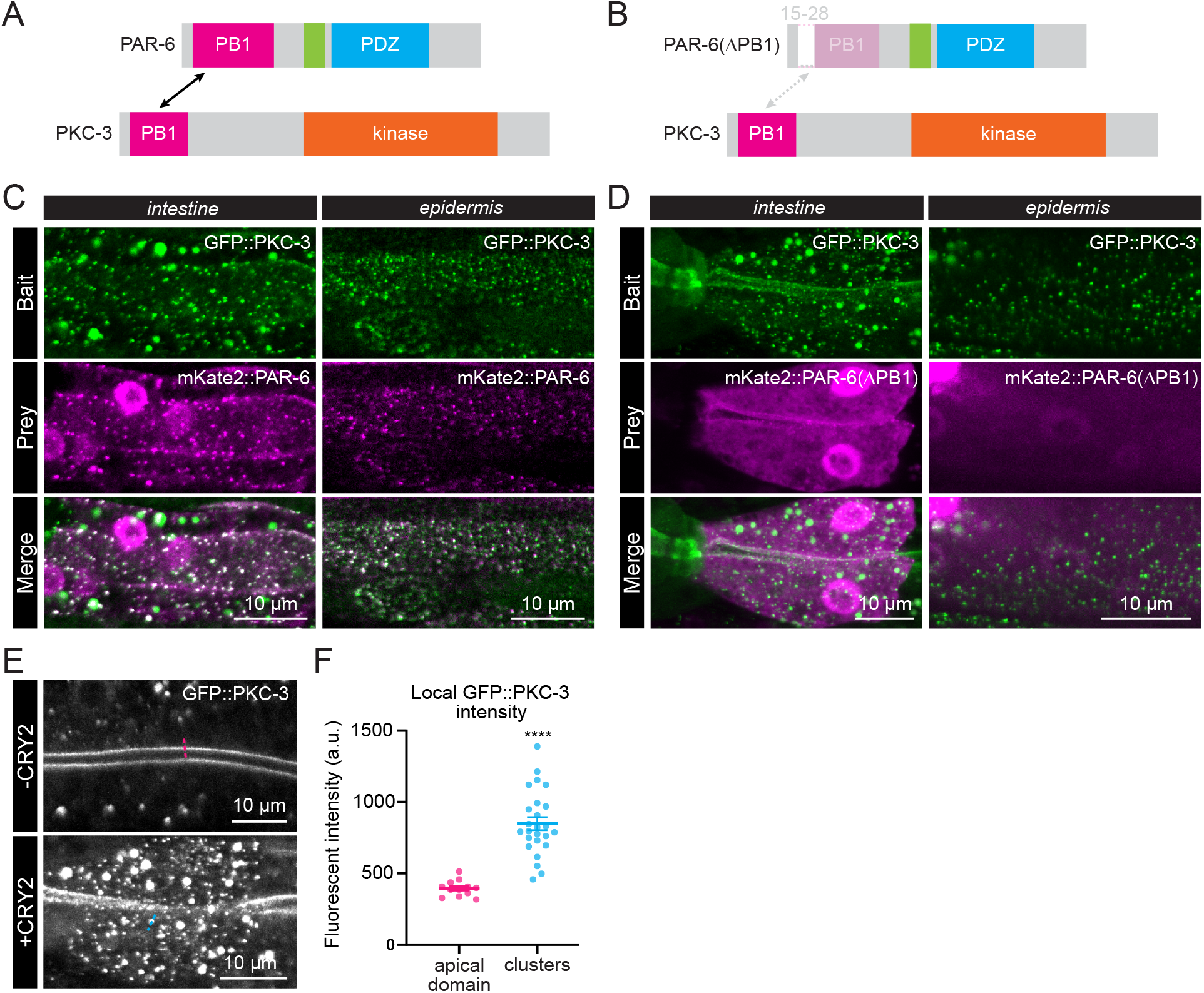
Interaction of PKC-3 with PAR-6 and an interaction defective PAR-6 variant assayed with CeLINC. (**A**) Schematic representation of the wild-type PKC-3 and PAR-6 proteins with labeled protein domains. Arrow indicates the PB1 domains that mediate the protein-protein interaction. (**B**) Diagram of PKC-3 and mutant PAR-6(ΔPB1) proteins with their domains. Dashed protein region in PAR-6 (aa 15–28) is deleted to abolish the interaction with PKC-3 (indicated by dashed arrow). (**C–D**) CeLINC interaction experiment for GFP::PKC-3 and mKate2::PAR-6 (C) or GFP::PKC-3 and mKate2::PAR-6(ΔPB1) (D) in the intestine and hypodermis. The GFP::PKC-3 bait protein is clustered by the nanobody fused to CRY2(olig). PAR-6 constructs were expressed from an extrachromosomal array. CeLINC constructs were expressed from the *rps-0* promoter. Overlapping clusters are white in the merged image. (**E**) Typical example of relative fluorescent intensity of GFP::PKC-3 at its native localization site at the apical membrane domain of the intestine (top) versus in clusters (bottom). (**F**) Quantification of GFP::PKC-3 fluorescent intensity. Data are represented as mean ± SEM and analyzed with unpaired t-test; **** = P < 0.0001. n = 12 measurements among 4 animals for the apical domain and n = 25 measurements among 5 animals for clusters.

The data above show that the CeLINC system can unambiguously identify and distinguish between a well characterized positive and negative protein-protein interaction pair in *C. elegans*. To further investigate the robustness of CeLINC, we tested three additional negative reference pairs: AID::GFP with either LGL-1::mScarlet or DLG-1::mCherry, and GFP::MAPH-1.1 with ERM-1::AID::mCherry. None of these combinations showed any co-clustering (Supplemental Data Figure S1). Overall, the CeLINC system can reliably identify a known protein interaction pair, and also does not show any false-positives in the negative reference pairs tested.

### Analysis of cortical cell polarity proteins with CeLINC assay

Next, we tested the assay with combinations of the apical polarity regulators PAR-3, PAR-6, and PKC-3 (aPKC) and the basolateral polarity regulators LGL-1, DLG-1, and LET-413. Each of these proteins was fluorescently tagged by CRISPR/Cas9 editing at the endogenous loci, preserving all aspects of normal localization and expression levels.

PAR-3, PAR-6, and PKC-3 are each localized to the apical membrane of intestinal cells and the junctional area of seam cells, among other epithelial tissues and cells types (Achilleos et al., 2010; Castiglioni et al., 2020; Li et al., 2010b; Nance et al., 2003; Welchman et al., 2007). PAR-3 is known to transiently, but not permanently, interact with the PAR-6/PKC-3 complex (Rodriguez et al., 2017). Testing the interaction of the PAR-6/PKC-3 complex with PAR-3 can determine whether CeLINC is able to detect more transient and dynamic protein interactions. We also tested the use of tissue-specific promoters directing the expression of the CRY2(olig) construct to the intestine and hypodermis, using the *elt-2* and *wrt-2* promoters, respectively. The CIBN-MP module was expressed from the *rps-0* general promoter. After light activation, the endogenously tagged PAR-6 and PKC-3 proteins showed extensive co-localization in cytoplasmic clusters in the intestine (Figure 4A) and the hypodermis (Figure 4B), as expected. Next, we tested the interaction of PAR-3 with the PAR-6/PKC-3 complex using GFP::PAR-3 and PAR-6::mCherry using the general promoter *rps-0* to express the CeLINC proteins. We identified co-localization between the two proteins, but fewer GFP clusters contained the mCherry signal than with the interaction between PAR-6 and PKC-3 (Figure 4C). This result is consistent with the transient and dynamic nature of the interaction. No co-clustering was observed in a negative control protein pair consisting of GPF::PKC-3 and DLG-1::mCherry, a basolateral protein not expected to interact with PKC-3 (Figure 5C).

**Figure 4.**
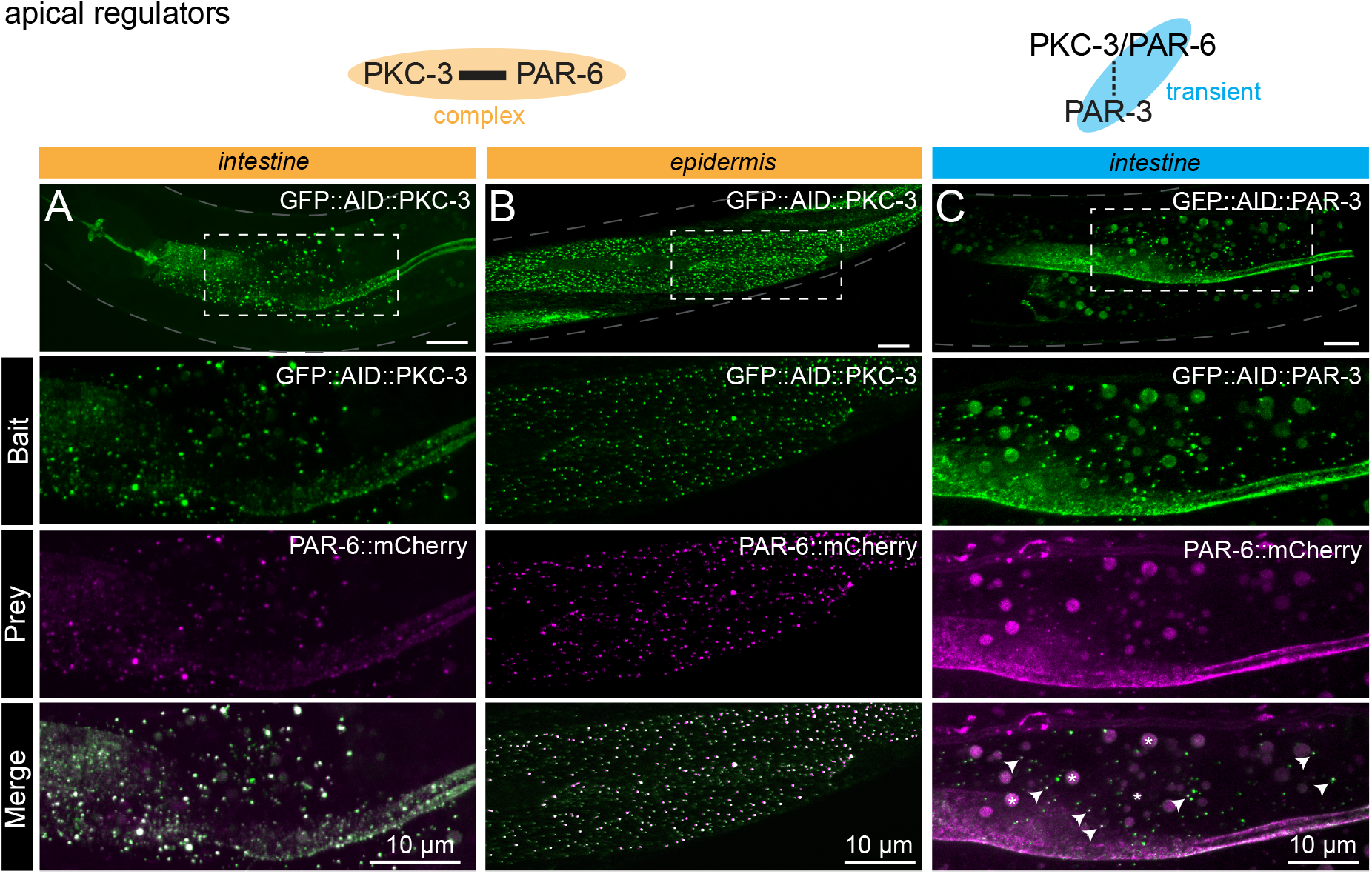
Interactions between apical cell polarity regulators assayed with CeLINC. (**A–B**) Interaction between GFP::AID::PKC-3 and PAR-6::mCherry using endogenously tagged alleles. The top panel shows the region of the worm examined and the area within the white dashed box is shown enlarged in the panels below. The CIBN-MP construct is expressed from the *rps-0* promoter. In (A), CRY2(olig)::VHH(GFP) is expressed from the *elt-2* promoter, while in (B), CRY2(olig)::VHH(GFP) is expressed from the *wrt-2* promoter to enable tissue-specific clustering. In all panels, the bait protein corresponds to the protein trapped by the CRY2-fused nanobody. (**C**) Interaction between GFP::AID::PAR-3 and PAR-6::mCherry. PAR-6 is not present in every GFP containing cluster, but PAR-6::mCherry clusters always overlap with PAR-3 clusters (white arrows indicate some of the co-clusters). Bigger round spheres in both the Bait and Prey channels correspond to autofluorescence from gut granules, which are marked with an asterisk in the merged image. CeLINC constructs are expressed from the *rps-0* promoter.

**Figure 5.**
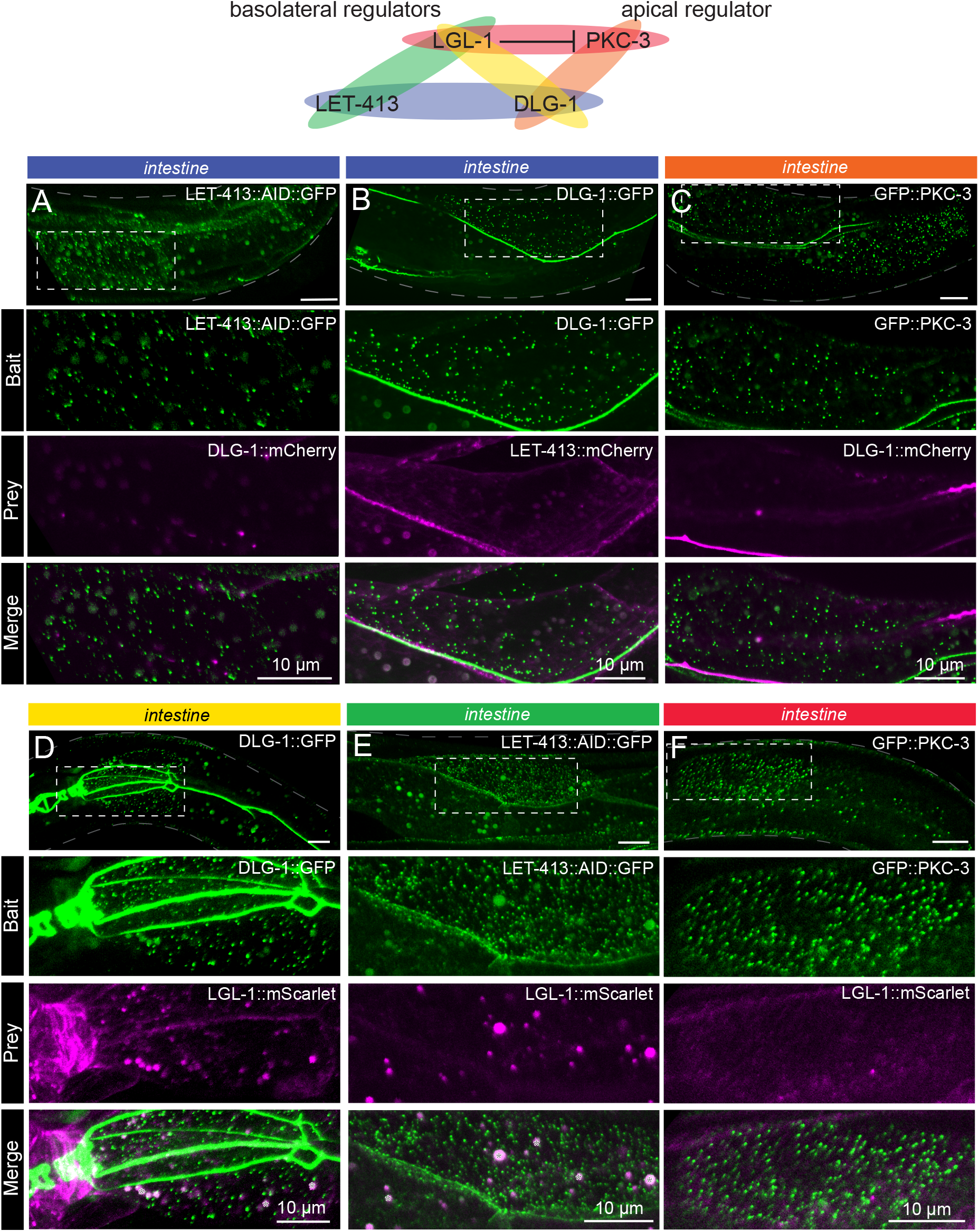
Basolateral cell polarity regulators assayed with CeLINC. (**A–F**) CeLINC experiments to investigate interactions among the basolateral cell polarity proteins LET-413, DLG-1, and LGL-1, and the apical protein PKC-3. The top panels show the region of the worm examined and the area within the white dashed box is shown enlarged in the panels below. Intestinal cells were analyzed, and all images represent maximum projections of a z-stack. The bait protein corresponds to the protein trapped by the CRY2 fused nanobody. All CeLINC constructs are expressed from the *rps-0* promoter. In (D–E), larger round spheres in both the Bait and Prey channels correspond to autofluorescence from gut granules, which are marked with an asterisk in the merged image. No physical interactions were detected in any of the basolateral protein pairs.

The Scribble module proteins LGL-1 (Lgl), DLG-1 (Dlg), and LET-413 (Scrib) play conserved roles in promoting basolateral domain identity, in part by antagonizing aPKC (Stephens et al., 2018; Wen and Zhang, 2018). In *Drosophila*, Lgl, Scrib, and Dlg are interdependent for their localization to the basolateral membrane in multiple tissues and act in a common basolateral polarity pathway (Bilder, 2000; Bilder et al., 2003; Khoury and Bilder, 2020; Su et al., 2012). However, unlike the apical polarity determinants, evidence for physical interactions between Scribble module members remains limited. In the synapses of Drosophila neuron, Dlg was also shown to indirectly associate with Scrib through the linker protein GUK-holder (Gukh) (Caria et al., 2018; Mathew et al., 2002). In mammalian cells, Lgl2 may interact with the guanylate kinase domain of Dlg4 (Zhu et al., 2014), as well as with the LAP unique region of Scrib (Choi et al., 2019; Kallay et al., 2006). Recently, the LINC system was used in *Drosophila* follicular epithelial cells to show that Dlg and Scribble interact via Scribble’s LRR domains (Ventura et al., 2020). However, the importance of these interactions for polarity establishment, and their conservation between organisms remains unclear.

Currently, there is no evidence in *C. elegans* that the basolateral proteins physically interact. Clarifying whether LGL-1, DLG-1, and LET-413 interact is important for understanding their function, and how their roles might differ between *C. elegans, Drosophila,* and mammals. LGL-1 and LET-413 have overlapping basolateral expression patterns in the intestine and the seam cells, whereas DLG-1 remains junctional in both cell types. We expressed CeLINC plasmids with general promoters and analyzed all combinations of the basolateral proteins with the CeLINC assay, but identified no co-clusters containing both signals (Figure 5A, B, D, E). Specifically, in contrast to the *Drosophila* follicular epithelium (Ventura et al., 2020), we found no co-clustering of LET-413 and DLG-1, despite testing for interaction in both orientations by using each protein separately as the bait protein targeted by the nanobody (Figure 5A, B). These results add to the body of evidence that these proteins do not belong to the same physical protein complex in *C. elegans* (Waaijers et al., 2016).

Finally, we investigate the interaction between LGL-1 and PKC-3. In the one-cell embryo, LGL-1 and PKC-3 engage in mutually inhibitory interactions to localize to opposing poles and promote cell polarity (Beatty et al., 2010; Hoege et al., 2010). In addition, depletion of PKC-3 in the epidermal seam cells causes apical invasion of LGL-1 (Castiglioni et al., 2020). Despite their localization to opposing membrane domains, PKC-3 and LGL-1 co-immunoprecipitate together (Hoege et al., 2010; Waaijers et al., 2016), which led to a model where LGL-1 associates with PAR-6/PKC-3 at the boundary of their respective domains, becomes phosphorylated by PKC-3, and subsequently dissociates from PAR-6/PKC-3 (Hoege et al., 2010). Using the CeLINC assay, we found that in the intestinal (Figure 5F) and hypodermal tissues (Supplementary Figure S2) there was no co-clustering between the proteins. Therefore, while CeLINC can detect some transient interactions, not all transient interactions are identified by the technique.

Overall, the CeLINC system was able to trap all of the cell polarity proteins tested into ectopic clusters within the cytoplasm of the intestinal cells, even DLG-1, which is localized to the cell junctions. We found no false-positives between any of the negative reference protein pairs tested, suggesting that the false-positive rate is low. In the positive reference pairs, we found that the interaction between PKC-3 and PAR-6 could be identified, and also prevented with a mutation of the PKC-3 binding site on PAR-6. Additionally, a more transient interaction, that of the PAR-6/PKC-3 complex and PAR-3, could also be identified with the assay.

### Generation of a CeLINC toolkit

To expand the potential use cases of CeLINC, we investigated the possibility of clustering proteins tagged with other fluorescent proteins than GFP. Since GFP and YFP share structural similarities, we tested the GFP nanobody against a YFP tagged protein, YFP::ACT-5. After injection with the CeLINC plasmids, we identified intestinal clusters in the YFP::ACT-5 strain, indicating YFP tagged proteins can also be used as bait proteins in the assay (Supplementary Figure S3). Next, we tested a nanobody targeting the mCherry protein (Yamagata and Sanes, 2018). We confirmed that CRY2(olig)::VHH(mCherry) was able to induce clustering of an mCherry tagged protein but not a GFP tagged protein (Figure 6A), and the mCherry nanobody could induce co-clustering of PAR-6 with PKC-3 (Figure 6B). The mCherry nanobody further expands the use of the system to allow for more proteins to be used as baits. Furthermore, if nanobodies against additional fluorescent proteins are developed they can be easily incorporated.

**Figure 6.**
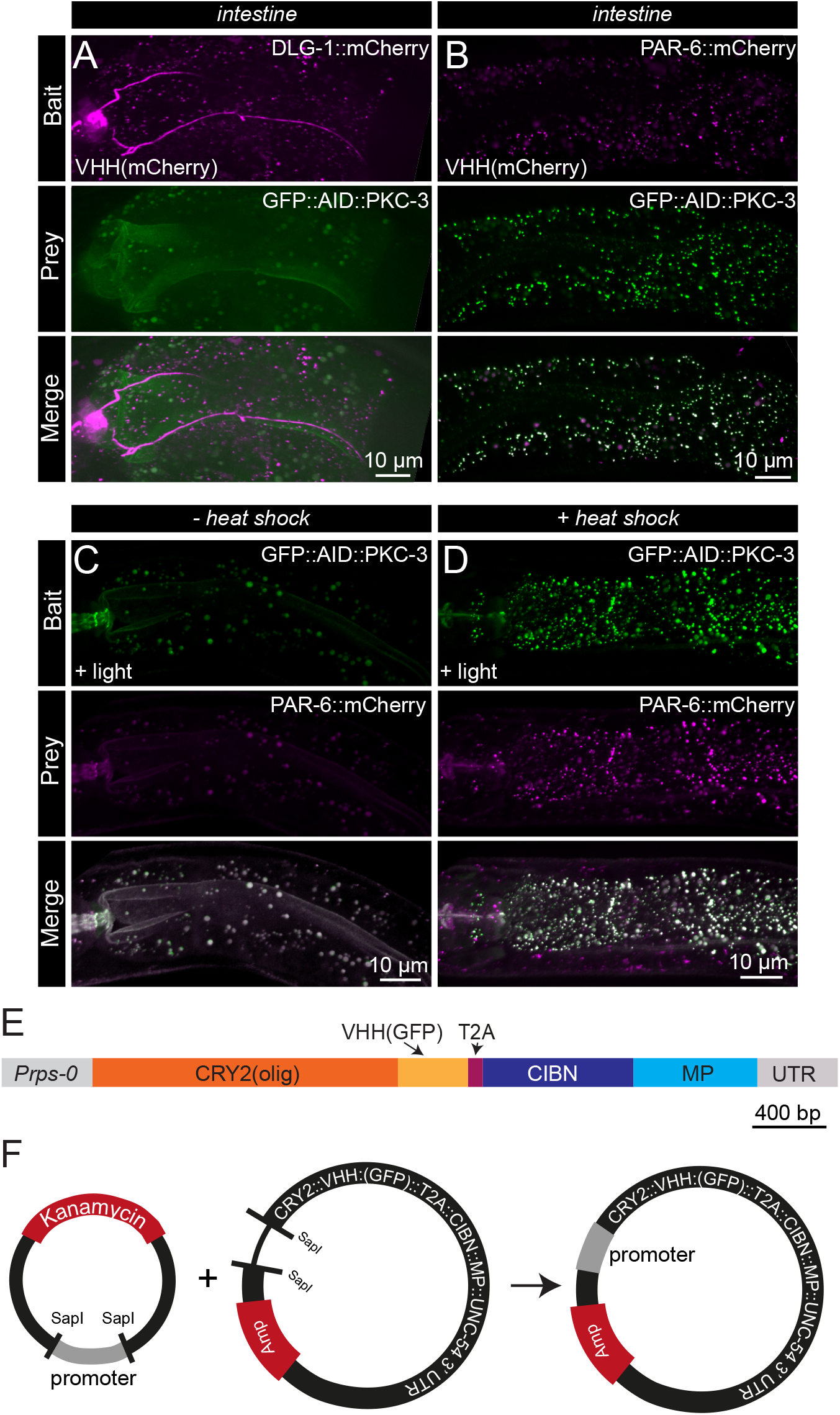
Expansion of CeLINC with additional modules and constructs. (**A–B**) Fluorescent maximum z-stack projections of negative (A) and positive (B) protein pairs assayed with CeLINC using an mCherry nanobody to trap the mCherry-tagged protein in clusters. In both protein pairs, the mCherry-tagged protein was recruited to clusters, but only in the positive protein pair (B) did the prey protein co-cluster. Intestinal cells were analyzed. (**C–D**) Interaction between PKC-3 and PAR-6 assessed using CRY2(olig) expressed from a heat shock promoter and the CIBN-MP protein from the *rps-0* promoter. Images are maximum z-stack projections of intestinal cells in animals kept at 20° C (C) or treated with a two-hour heat shock at 30° C (D). Larger round spheres in both the Bait and Prey channels in the non-heat shocked animals correspond to autofluorescence from gut granules. (**E**) Diagram of the coding region of the *Prps-0*::CRY2(olig)-T2A-CIBN-MP plasmid (pJRK260), which allows for both CeLINC proteins to be translated from the same mRNA molecule. (**F**) Schematic of the CRY2-T2A-CIBN-MP destination plasmid (pJRK261) which contains an empty promoter module. The desired promoter donor plasmid can be combined and assembled with the SapTrap method in one-step to generate a functional and complete CeLINC plasmid.

In order to be able to easily adapt or change the components of CeLINC, we have made the plasmid cloning system modular with the use of the SapTrap plasmid assembly method (Schwartz and Jorgensen, 2016). In this way, different promoters, nanobodies, CRY2 variants, or 3’ UTRs can be combined with previously generated donor plasmids and assembled into the final destination vector (Supplementary Data Table S2 and Table S4). We have already generated a series of CRY2(olig) plasmids with more specific promoters, such as tissue specific promoters for the intestine (*elt-2*), the hypodermis (*wrt-2*) (Figure 4A, B), and a heat-shock promoter (*hsp-16.48*) (Figure 6C, D). The heat-shock promoter might be useful for some tagged-protein combinations, since even in the dark, nanobodies will still be targeted to GFP or mCherry proteins, which could cause unintended effects for certain types of proteins (though no problems have been identified with the proteins tested thus far). We kept CIBN-MP with the *rps-0* promoter since it is the more “passive” element of the CeLINC system. We have also used the T2A system that allows two peptides to be produced from the same mRNA (Ahier and Jarriault, 2014), and have created a single plasmid encoding both the CRY2(olig)::VHH(GFP) nanobody element and the CIBN-MP element from the *rps-0* promoter (Figure 6E). Finally, we generated a version of this T2A based plasmid with an empty promoter module with flanking *SapI* sites so that any promoter can be swapped in with a single donor plasmid and reaction (Figure 6F). This toolkit of plasmids will allow any particular tissue type or cell type to be targeted with the CeLINC system and increases the types of proteins that can be used as bait proteins in the assay.

## Discussion

We have adapted and tested the light-induced co-clustering assay (LINC), for use in *C. elegans*. We have shown that in *C. elegans*, expressed CRY2(olig) is activated by blue light, and efficiently clusters in multiple tissues, cell types, and cellular compartments. When we compared the interaction of PKC-3 with a wild-type PAR-6 protein and an interaction-defective mutant, we saw a clear difference in the co-clustering of PAR-6. We tested additional negative reference protein pairs and no false positives were detected among them. Testing for physical interactions among apical and basolateral cell polarity regulators using the system also did not identify false-positive interactions, and successfully recapitulated the known interaction between PAR-3 and PAR-6. Finally, we developed a toolkit of plasmids to enable flexibility and adaptability of the system for future uses.

The siphoning of proteins from their endogenous location to clusters or to ectopic locations within the cell could cause gain or loss-of-function phenotypes, as the original purpose of the LARIAT system was to disrupt protein function (Lee et al., 2014). We saw no apparent lethality or toxicity associated with CRY2/CIBN-MP expression in any of the strains generated. Therefore, the CeLINC system appears to have little detrimental effect on the fitness of animals, even when clustering cell polarity proteins that are essential for animal development. However, throughout our experiments, we limited the amount of light exposed to the animals. Longer exposure of the animals to blue or bright light conditions could cause developmental or cellular phenotypes. Moreover, since every protein may behave differently or have different thresholds for a “knock sideways”-like inhibition, other protein combinations or bait proteins may still show unexpected phenotypes or effects. Finally, while we observed regularly sized and spaced cluster formation, it is possible that CRY2 clusters may form in different shapes or sizes with the use of different bait proteins, depending on what interactions the bait protein normally engages in. For example, filamentous proteins or proteins strongly associated with a membrane may produce clusters that are larger or more amorphous in shape.

One consideration for the proper interpretation of the assay is protein mobility or accessibility. Some proteins could be resistant to clustering or less likely to form ectopic clusters than other proteins. Proteins unable to mis-localize from their endogenous localization would produce a false-negative result if used as a prey protein in the assay. For example, fewer clusters formed in the LET-413 (Scribble) strains when used as a bait, suggesting that LET-413 protein is either tightly bound or highly integrated to its endogenous location in the cell. However, since each extrachromosomal array in our experiments resulted from a distinct injection, differences in clustering ability could also be attributed to differing levels of CeLINC construct expression. Similar to other PPI assays, a negative result needs to be taken with caution. However, positive results from the assay are likely to be highly significant, since no false-positives were identified in any negative protein pair that was tested, even among basolateral proteins that co-localize. Additionally, the assay is limited by the intensity of the particular fluorophore and expression level of the target proteins. While one advantage is that the clusters concentrate the signal in a small area, giving a bright focused spot (Figure 3E), some lowly expressed proteins are still barely visible. For example, red fluorescent proteins tend to have a reduced signal compared to GFP, so weakly expressing proteins might be prioritized to be tagged with GFP. Another way to overcome low expression of target proteins is to use overexpression constructs. The use of transgenes can also circumvent detrimental phenotypes and is necessary to test proteins carrying non-viable mutations, as was the case with *par-6(ΔPB1)*. Finally, like many protein-protein interaction assays, CeLINC cannot distinguish between indirect and direct protein-protein interactions.

While light activation of CRY2 formation is useful for temporal control of cluster formation, for the purposes of the CeLINC assay, blue light activation and cluster formation during the microscopy session is not necessarily needed. Pre-activation of the clusters during mounting the animals on slides increased the throughput of the assay, as more animals could be imaged per slide, and animals exhibiting clusters could be more rapidly identified. In our experiments, we easily identified animals and cells expressing the CeLINC constructs because GFP or mCherry clusters were distributed in the cytoplasm of the cells, a completely different location than the normal localization of the tagged proteins. Additionally, due to mosaicism of the extrachromosomal array, surrounding cells where no clusters were observed could often be used as negative controls within the same animal. This ensured we were not analyzing cluster-like aggregates, gut granules that are auto fluorescent, or endogenous localization of the proteins. Nevertheless, light activation of CRY2 on the microscope might be useful for certain tagged proteins that are already prone to aggregation or exhibit an endogenous localization pattern that might easily be confused with CRY2 mediated clusters.

The CeLINC assay is mainly a binary assay, answering whether two particular proteins do or do not interact. However, with the use of particle analysis software or tools, more quantitative measurements could be made, such as the degree of clusters that contain a fluorescent signal of the prey protein. However, caution should be given when comparing different combinations of proteins, since their degree of co-clustering may be influenced by other factors than purely their physical association.

Compared to other PPI assays available for use in *C. elegans*, CeLINC uses relatively few special reagents, is rapid, and is straight-forward to interpret. Many proteins under study already have fluorescently tagged alleles available, and no further modifications need to be made for use in the CeLINC system. While we mainly use proteins tagged with mCherry, proteins tagged with YFP, mScarlet, mKate2, or BFP could be used with the system. With the recent development of CRISPR/Cas9 editing techniques and split protein fluorescent based systems, such as sfGFP_11_ and Split-mScarlet_11_ (Goudeau et al., 2021), where only a small fragment of the fluorescent protein needs to be integrated in a genetic background expressing the complementary fragment, fluorescent protein tags can be made with relative ease. Overall, the CeLINC system is a powerful technique to study protein-protein interactions that can utilize many existing strains and produces a clear result with commonly available equipment in any *C. elegans* laboratory.

## Materials and Methods

### Plasmid cloning

Plasmid names and descriptions are available in Supplementary Data Table S2. Primer information is in Supplementary Data Table S3. SapTrap donor plasmid overhangs and assembly information are found in Supplementary Data Table S4. Annotated GenBank files of the plasmids are available in Supplementary File 2. Plasmids will be made available at Addgene (https://www.addgene.org/).

Sequences for CRY2(olig) (Taslimi et al., 2014), CIBN(1–170) (Lee et al., 2014), mCherry nanobody (RANbody2 mCherry nanobody variant) (Yamagata and Sanes, 2018), and MP (Lee et al., 2014) were codon-optimized using the *C. elegans* codon adapter tool (Redemann et al., 2011) and ordered as gBlocks Gene Fragments (IDT) with appropriate *SapI* restriction sites and overhangs flanking the sequences. The MP, CIBN, and CRY2(olig) sequences each contain one synthetic intron. A 3xFLAG tag and linker segment was added to the C-terminus of the CIBN sequence. The CRY2 variant used, CRY2(olig), contains an E490G mutation to increase clustering ability (Taslimi et al., 2014). The GFP nanobody (VHH(GFP)) sequence was PCR amplified from plasmid pVP130 (Vaart et al., 2020), which was codon-optimized from the original sequence (Wang et al., 2017). The *Pwrt-2* promotor was amplified from plasmid pRS177, the *Pelt-2* promoter was amplified from plasmid pSMR12, the *Phsp-16.48* promoter from plasmid pJRK83, and the *par-6* coding sequence from pJRK11. *SapI* restriction sites and overhangs for SapTrap assembly were included in the primers used for amplification. PCR amplicons and gBlock fragments were phosphorylated and cloned blunt-ended into the plasmid backbone pHSG298 digested with *Eco53KI*. Donor plasmids were combined as shown in Supplementary Data Table S4. The SapTrap assembly method (Schwartz and Jorgensen, 2016) was used to assemble donor plasmids to generate the final expression plasmids used for injections. Donor vector mixes were then combined with pMLS257 predigested and linearized with *SapI*. pMLS257 was a gift from Erik Jorgensen (Addgene plasmid # 73716) (Schwartz and Jorgensen, 2016). The plasmid pJRK86 for general AID::GFP expression was made by combining the donor plasmids pJRK1, pDD397, pJRK245, and pJRK150.

The co-injection plasmid pJRK248 (*Prps-0::HygR::unc-54 3’UTR; Psqt-1::sqt-1::sqt-1 3’* UTR) used in this study contains the dominant markers HygR (Hygromycin resistance) and a *sqt-1* mutation (conferring Roller phenotype), and is a derivative of the plasmid pDD382. pDD382 was a gift from Bob Goldstein (Addgene plasmid # 91830). To create pJRK248, two PCR fragments were generated that excluded the heat-shock Cre module of pDD382, and then reassembled via Gibson assembly. The plasmid pJRK259 (*Prps-0::mKate2::par-6(ΔPB1*)::unc*-54 3’UTR*) was made by Gibson assembly from the plasmid pJRK258 (*Prps-0::mKate2::par-6::unc-54 3’UTR*) using two PCR fragments that excluded the amino acids 15–28 of the *par-6* coding region. Plasmids pJRK260 and pJRK261 were made by Gibson assembly. Four fragments were used to assemble pJRK260: three fragments originating from PCR products (pJRK136 and pJRK138 were used as templates) and the T2A segment consisting of a gBlock Gene Fragment (IDT). Three fragments were used to assemble pJRK261: two fragments originating from PCR products (pJRK260 was used as the template) and the empty promoter/SapI region consisting of a gBlock Gene Fragment.

### Strains and generation of extrachromosomal array strains

Strains are available upon request. The complete list of genotypes and strains used is in Supplementary Data Table S1. N2 was used as the wild-type strain. The *pkc-3(it309[gfp::pkc-3])* allele was linked to the *dpy-10(cn64)* allele in strain FT1991 and precluded efficient injection. Therefore, the strain was first crossed to N2 to isolate the *lgl-1(xn103[lgl-1::zf1::mScarlet])* allele, and then was crossed to *pkc-3(it309[gfp::pkc-3])* to generate the strain BOX757.

Fluorescently tagged strains were first generated by crossing and then CeLINC plasmids were injected into the gonads of young adult worms to form extrachromosomal arrays. Injection mixes were made with the CRY2(olig)::nanobody and CIBN-MP plasmids at a concentration of 10 ng/μL, and the co-injection plasmid pJRK248 at 20 ng/μL. Strains with *mKate2::par-6* or *mKate2::par-6(ΔPB1)* were at a concentration of 10 ng/μL. Negative reference pair strains in Supplemental Figure S1A, B had injection mixes with 10 ng/μL of pJRK86 (AID::GFP). Lambda DNA (Thermo Scientific) was used to bring the total concentration of DNA to 95 ng/μL. Plasmids were isolated from bacteria using the HQ PureLink Mini Plasmid Purification Kit (Invitrogen) with an extra wash step. After injection, transgenic animals were identified by a Rol phenotype and/ or resistance to hygromycin. For hygromycin selection, 300–400 μL of 5 mg/ml hygromycin B (Foremedium Ltd) dissolved in water was added to the plates 2–3 days after growth on the plate was established.

### Animal handling

Animals were grown on standard nematode growth medium (NGM) agar plates at 20° seeded with OP50. Hermaphrodites in the L2–L4 larval stage were used for imaging. Animals grown in the ambient light condition were grown in the dark but mounted under a binocular microscope with normal white light illumination. Animals grown in the complete darkness condition were grown in the dark and mounted for microscopy in dark conditions with only the use of a green or red light in the room. The slide was then transported in aluminum foil to the spinning disk microscope. The animals were focused and moved into position on the microscope using the dimmest possible setting of a white light. For the heat-shock experiment in Figure 6C and 6D, worms were either kept at 20° (−heat-shock) or 30° (+heat-shock) for two hours, and then imaged.

### Imaging and blue light activation

Imaging was performed by mounting larvae on a 5% agarose pad in 1 mM levamisole solution in M9 buffer to induce paralysis. Images were taken with a Nikon Ti-U microscope driven by MetaMorph Microscopy Automation and Image Analysis Software (Molecular Devices) and equipped with a Yokogawa CSU-X1-M1 confocal head and an Andor iXon DU-885 camera, using a 60× 1.4 NA objective and with 0.25 μm z-step intervals. Exposure settings were customized for each fluorescently tagged protein, due to wide variations in expression levels and signal intensities. To activate cluster formation with blue light from starting dark conditions, as in Figure 2, z-stacks were taken of the sample with the blue laser with 300ms exposure, 50% laser power, and 50–80 z-stacks (depending on the sample depth). For Figures 3–6, animals were mounted with white light therefore clusters were pre-activated before imaging. Animals were then directly imaged, and z-stacks were obtained with both green and blue lasers. Images were analyzed and processed with ImageJ/FIJI (Schindelin et al., 2012). Photoshop was used to non-destructively prepare images and Adobe Illustrator was used for figure preparation.

### Half-life experiment & cluster quantification

For the experiment in Figure 2B and C, animals were grown and mounted in complete darkness. The animal was imaged first with the green laser to determine the baseline mKate2 signal (pre blue light). Next, the clusters were activated by imaging the z-stack with both green and blue lasers (300ms exposure, 50% laser power, and 89–105 z-stacks), corresponding to the 0 min time-point. After activation, the animal was imaged in 5-minute intervals for 25 minutes with only the green laser. Maximum projections of the z-stack were generated at each time point, and ComDet 0.5.5 plugin for ImageJ (https://github.com/ekatrukha/ComDet) was used to detect and count the number of clusters. The parameters used were: “include larger particles”-true, “segment larger particles”-false. In animals one and two, approximate particle size was set to 3.0, and intensity threshold (in SD) was set to 25. For animal three in the experiment, “include larger particles” was set to false, the approximate particle size was 2.0, and the intensity threshold (in SD) was increased to 35. Cluster numbers were normalized by subtracting the number of clusters identified in the pre-blue-light timepoint from the number of clusters in the following time-points to have a baseline corresponding to zero clusters. The maximum number of clusters identified in each sample was then used to determine the fraction of maximum clusters at every time point. The half-life was determined by solving the equation

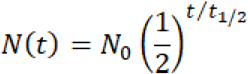

for *t_1/2_*, where *N_0_* is the initial quantity, *N(t)* is the remaining quantity after time *t*, and *t_1/2_* is the half-life. For Figure 1E and F, the following settings were used for the ComDet plugin: “include larger particles”-true, “segment larger particles”-false, approximate particle size was set to 2.0, and intensity threshold (in SD) was set to 5.

## Acknowledgements and Funding

Thanks to Vincent Portegijs and Ruben Schmidt for plasmids, and Amalia Riga, Victoria Castiglioni, Jorian Sepers, and Helena Pires for strains. We thank members of the S. van den Heuvel, S. Ruijtenberg, and M. Boxem groups for helpful discussions. Some strains were provided by the Caenorhabditis Genetics Center (CGC), which is funded by NIH Office of Research Infrastructure Programs (P40 OD010440). This work was supported by a OCENW. XS3.087 grant to J. Kroll and a NWO-VICI 016.VICI.170.165 grant to M. Boxem.

## Supplementary Data

**Figure S1.**
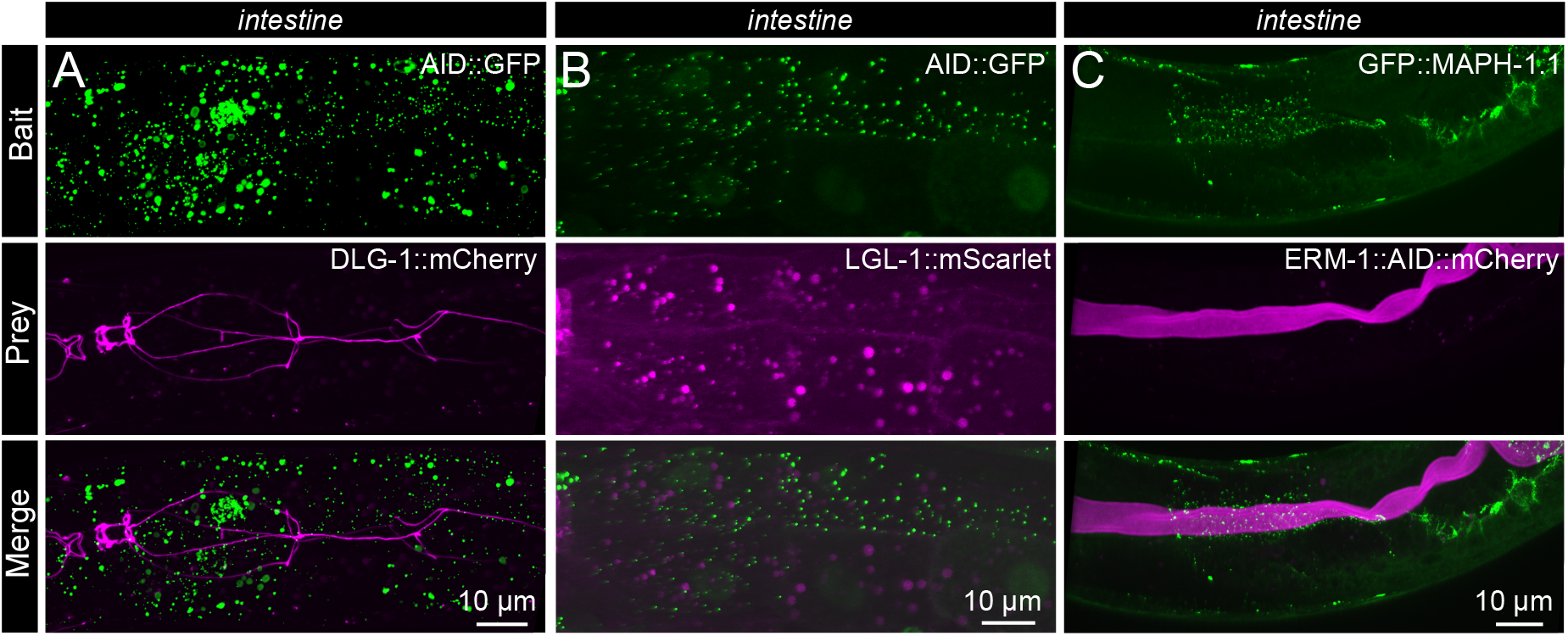
Additional negative protein interaction pairs tested with CeLINC. (**A–C**) Additional negative protein pairs tested with CeLINC. In (A) and (B), the bait protein used was AID::GFP expressed from a general promoter (*rps-0*), with DLG-1 and LGL-1 endogenously tagged with mCherry or mScarlet as the prey proteins. In (C) the bait protein was the microtubule binding protein MAPH-1.1 endogenously tagged with GFP, and the prey protein ERM-1 endogenously tagged with AID::mCherry. The CeLINC constructs were expressed from the *rps-0* promoter. Images represent maximum projections of a z-stack. No co-clusters were observed in the intestinal cells of these animals.

**Figure S2.**
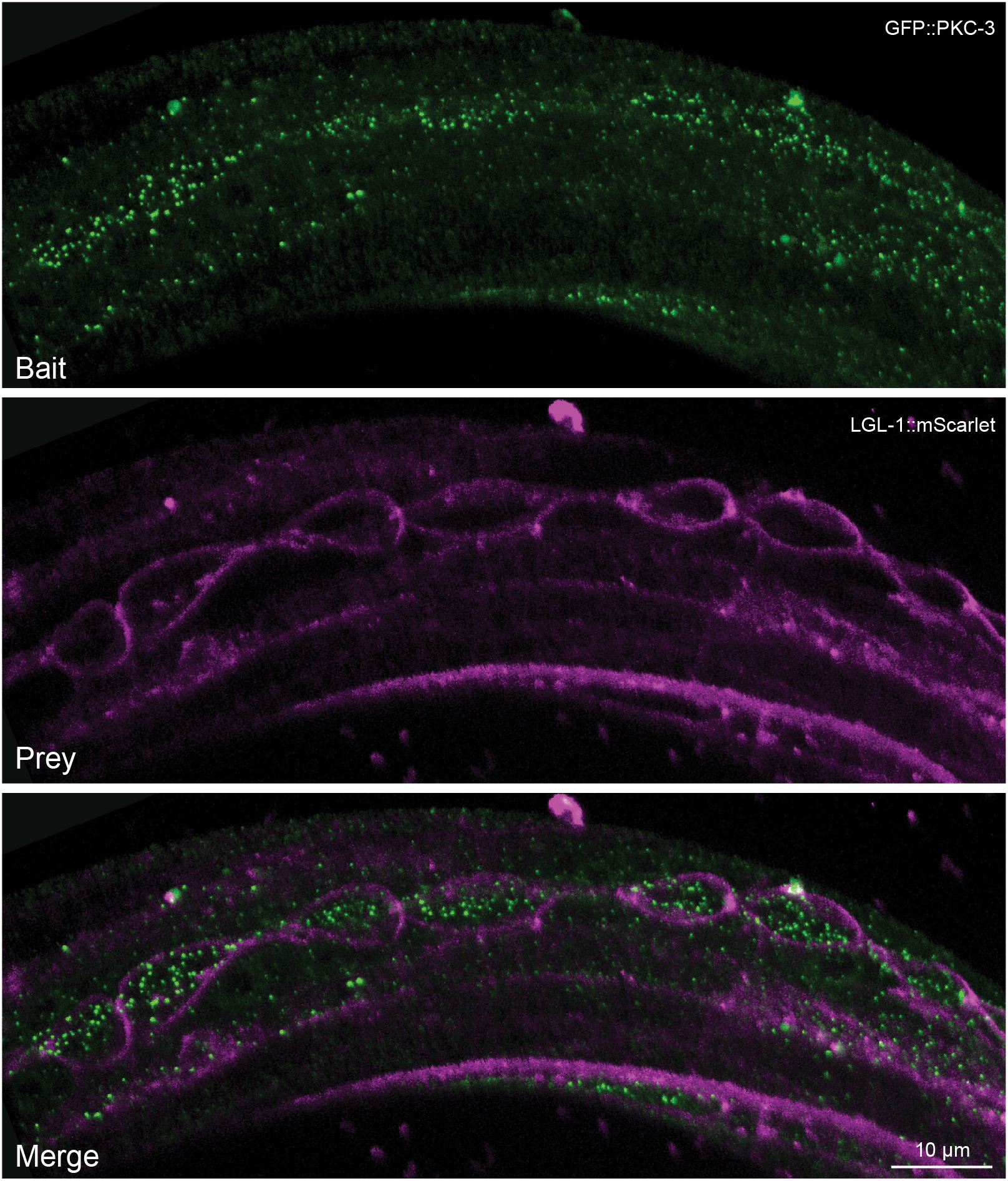
CeLINC assay for PKC-3 and LGL-1 in the hypodermis. CeLINC assay between endogenously tagged GFP::PKC-3 as bait and LGL-1::mScarlet as prey. The CeLINC constructs were expressed from the general *rps-0* promoter. Images represent maximum projections of a z-stack. No co-clusters were observed in the seam cells or hypodermis, despite LGL-1 being a target of the PKC-3 kinase.

**Figure S3.**
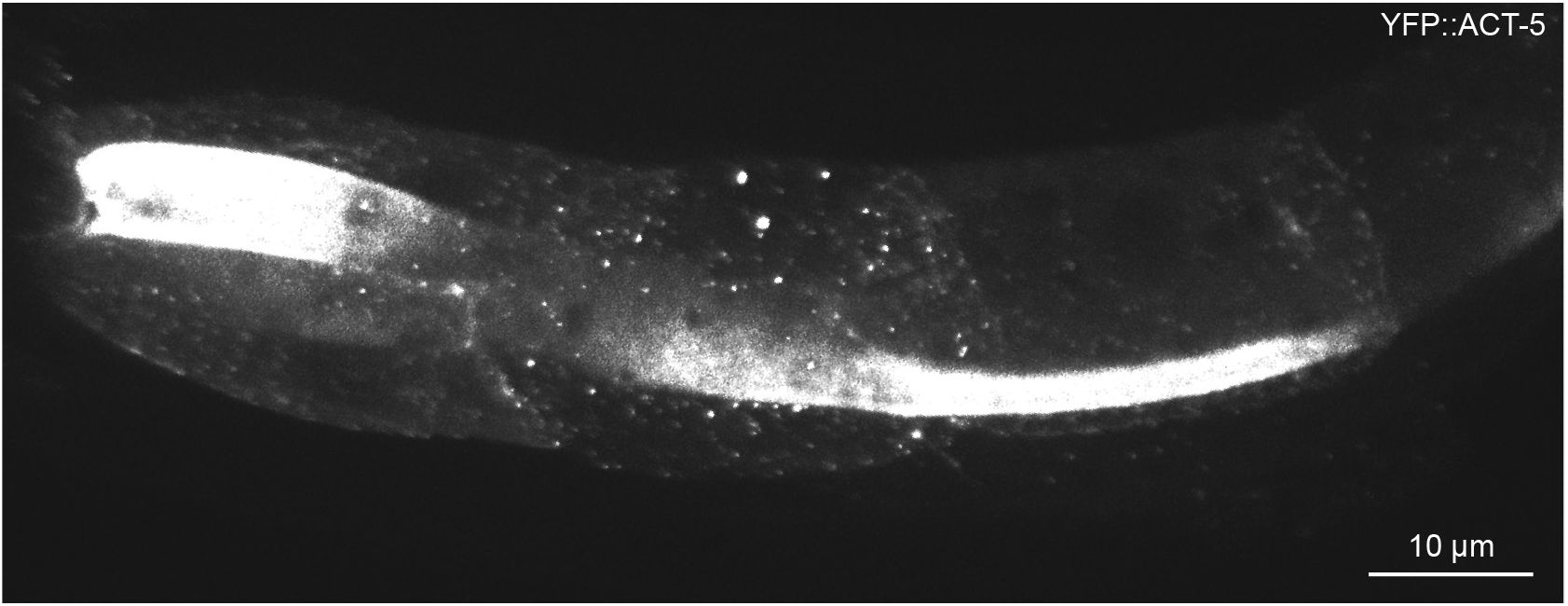
YFP can be captured into clusters by the GFP nanobody. Image of the intestine of an animal with a YFP::ACT-5 transgene expressing mKate2::CRY2(olig)::VHH(GFP) and CIBN-MP plasmids from a general promoter (*rps-0*). Due to their structural similarity, the YFP tagged protein was captured into clusters by the GFP nanobody.

**Table S1.** *C. elegans* strains.

| Name | Genotype | Reference |
| --- | --- | --- |
| **BOX249** | *let-413(mib29[let-413::mCherry-LoxP]) V; dlg-1(mib22[dlg-1::eGFP-LoxP]) X* | This study |
| **BOX451** | *let-413(mib32[let-413::AID::eGFP-LoxP]) V; dlg-1(mib23[dlg-1::mCherry-LoxP]) X* | This study |
| **BOX484** | *par-6(mib25[par-6::mCherry-LoxP]) I; pkc-3(mib78[eGFP-LoxP::AID::pkc-3]) II; mibls49[Pwrt-2::TIR-1::tagBFP2-Lox511::tbb-2-3'UTR, IV:5014740-5014802 (cxTi10882 site))] IV* | (Castiglioni et al., 2020) |
| **BOX486** | *par-6(mib25[par-6::mCherry-LoxP]) I; par-3(mib68[eGFP-Lox-2272::AID::par-3b+eGFP(noIntrons)-LoxP::AID::par-3g]) III; mibls49[Pwrt-2::TIR-1::tagBFP2-Lox511::tbb-2-3'UTR, IV:5014740-5014802 (cxTi10882 site))] IV* | (Castiglioni et al., 2020) |
| **BOX493** | *pkc-3(mib78[eGFP-LoxP::AID::pkc-3]) II; mibls49[Pwrt-2::TIR-1::tagBFP2-Lox511::tbb-2-3'UTR, IV:5014740-5014802 (cxTi10882 site))] IV; dlg-1(mib23[dlg-1::mCherry-LoxP]) X* | (Castiglioni et al., 2020) |
| **FT1991** | *dpy-10(cn64) pkc-3(it309[GFP::pkc-3]) II; lgl-1(xn103[lgl-1::zf1::mScarlet]) X* | (Montoyo-Rosario et al., 2020) |
| **BOX740** | *lgl-1(xn103[lgl-1::zf1::mScarlet]) X* | This study |
| **BOX756** | *lgl-1(xn103[lgl-1::zf1::mScarlet]) dlg-1(mib22[dlg-1::eGFP-LoxP]) X* | This study |
| **BOX757** | *pkc-3(it309[GFP::pkc-3]) II; lgl-1(xn103[lgl-1::zf1::mScarlet]) X* | This study |
| **BOX758** | *let-413(mib81[GFP::LoxP::AID::let-413] V; lgl-1(xn103[lgl-1::zf1::mScarlet]) X* | This study |
| **KK1228** | *pkc-3(it309[GFP::pkc-3]) II* |  |
| **BOX235** | *dlg-1(mib23[dlg-1::mCherry-LoxP]) X* | This study |
| **BOX377** | *erm-1(mib40[erm-1::AID::mCherry]) maph-1.1(mib12[GFP::maph-1.1]) I* | This study |
| **JM125** | *cals[Pges-1::YFP::ACT-5]* | (Bossinger et al., 2004) |
| **BOX742** | *mibEx255[Prps-0::mKate2-CRY2olig-VHH(GFP)::unc-54 3'UTR; Prps-0::CIBN-3xFLAG-MP::unc-54 3'UTR; pJRK248, $\lambda$ DNA]* | This study |
| **BOX743** | *mibEx254[Prps-0::mKate2-CRY2olig-VHH(GFP)::unc-54 3'UTR; pJRK248, $\lambda$ DNA]* | This study |
| **BOX759** | *par-6(mib25[par-6::mCherry-LoxP]) I; pkc-3(mib78[eGFP-LoxP::AID::pkc-3]) II; mibls49[Pwrt-2::TIR-1::tagBFP2-Lox511::tbb-2-3'UTR, IV:5014740-5014802 (cxTi10882 site))] IV; mibEx259[Pwrt-2::CRY2olig-VHH(GFP)::unc-54 3'UTR; Prps-0::CIBN-3xFLAG-MP::unc-54 3'UTR; pJRK248, $\lambda$ DNA]* | This study |
| **BOX760** | *par-6(mib25[par-6::mCherry-LoxP]) I; pkc-3(mib78[eGFP-LoxP::AID::pkc-3]) II; mibls49[Pwrt-2::TIR-1::tagBFP2-Lox511::tbb-2-3'UTR, IV:5014740-5014802 (cxTi10882 site))] IV; mibEx260[Pelt-2::CRY2olig-VHH(GFP)::unc-54 3'UTR; Prps-0::CIBN-3xFLAG-MP::unc-54 3'UTR; pJRK248, $\lambda$ DNA]* | This study |
| **BOX761** | *par-6(mib25[par-6::mCherry-LoxP]) I; par-3(mib68[eGFP-Lox-2272::AID::par-3b+eGFP(noIntrons)-LoxP::AID::par-3g]) III; mibls49[Pwrt-2::TIR-1::tagBFP2-Lox511::tbb-2-3'UTR, IV:5014740-5014802 (cxTi10882 site))] IV mibEx256[Prps-0::CRY2olig-VHH(GFP)::unc-54 3'UTR; Prps-0::CIBN-3xFLAG-MP::unc-54 3'UTR; pJRK248, λ DNA]* | This study |
| **BOX762** | *let-413(mib29[let-413::mCherry-LoxP]) V; dlg-1(mib22[dlg-1::eGFP-LoxP]) X; mibEx256[Prps-0::CRY2olig-VHH(GFP)::unc-54 3'UTR; Prps-0::CIBN-3xFLAG-MP::unc-54 3'UTR; pJRK248, λ DNA]* | This study |
| **BOX763** | *let-413(mib32[let-413::AID::eGFP-LoxP]) V; dlg-1(mib23[dlg-1::mCherry-LoxP]) X; mibEx256[Prps-0::CRY2olig-VHH(GFP)::unc-54 3'UTR; Prps-0::CIBN-3xFLAG-MP::unc-54 3'UTR; pJRK248, λ DNA]* | This study |
| **BOX764** | *pkc-3(mib78[eGFP-LoxP::AID::pkc-3]) II; mibls49[Pwrt-2::TIR-1::tagBFP2-Lox511::tbb-2-3'UTR, IV:5014740-5014802 (cxTi10882 site))] IV; dlg-1(mib23[dlg-1::mCherry-LoxP]) X; mibEx256[Prps-0::CRY2olig-VH-H(GFP)::unc-54 3'UTR; Prps-0::CIBN-3xFLAG-MP::unc-54 3'UTR; pJRK248, λ DNA]* | This study |
| **BOX765** | *par-6(mib25[par-6::mCherry-LoxP]) I; pkc-3(mib78[eGFP-LoxP::AID::pkc-3]) II; mibls49[Pwrt-2::TIR-1::tagBFP2-Lox-511::tbb-2-3'UTR, IV:5014740-5014802 (cxTi10882 site))] IV; mibEx261[Phsp-16.48::CRY2olig-VHH(GFP)::unc-54 3'UTR; Prps-0::CIBN-3xFLAG-MP::unc-54 3'UTR; pJRK248, λ DNA]* | This study |
| **BOX766** | *par-6(mib25[par-6::mCherry-LoxP]) I; pkc-3(mib78[eGFP-LoxP::AID::pkc-3]) II; mibls49[Pwrt-2::TIR-1::tagBFP2-Lox511::tbb-2-3'UTR, IV:5014740-5014802 (cxTi10882 site))] IV; mibEx258[Prps-0::CRY2olig-VHH(mCherry)::unc-54 3'UTR; Prps-0::CIBN-3xFLAG-MP::unc-54 3'UTR; pJRK248, λ DNA]* | This study |
| **BOX767** | *lgl-1(xn103[lgl-1::zf1::mScarlet]) dlg-1(mib22[dlg-1::eGFP-LoxP]) X; mibEx256[Prps-0::CRY2olig-VHH(GFP)::unc-54 3'UTR; Prps-0::CIB1-3xFLAG-MP::unc-54 3'UTR; pJRK248, λ DNA]* | This study |
| **BOX768** | *pkc-3(it309[GFP::pkc-3]) II; lgl-1(xn103[lgl-1::zf1::mScarlet]) X; mibEx256[Prps-0::CRY2olig-VHH(GFP)::unc-54 3'UTR; Prps-0::CIB1-3xFLAG-MP::unc-54 3'UTR; pJRK248, λ DNA]* | This study |
| **BOX769** | *let-413(mib81[gfp::LoxP::AID::let-413]) V; lgl-1(xn103[lgl-1::zf1::mScarlet]) X; mibEx256[Prps-0::CRY2olig-VHH(GFP)::unc-54 3'UTR; Prps-0::CIB1-3xFLAG-MP::unc-54 3'UTR; pJRK248, λ DNA]* | This study |
| **BOX770** | *pkc-3(it309[GFP::pkc-3]) II; mibEx267[Prps-0::mKate2-par-6::unc-54 3'UTR; Prps-0::CRY2olig-VHH(GFP)::unc-54 3'UTR; Prps-0::CIB1-3xFLAG-MP::unc-54 3'UTR; pJRK248, λ DNA]* | This study |
| **BOX771** | *pkc-3(it309[GFP::pkc-3]) II; mibEx268[Prps-0::mKate2-par-6(ΔP-B1)::unc-54 3'UTR; Prps-0::CRY2olig-VHH(GFP)::unc-54 3'UTR; Prps-0::CIB1-3xFLAG-MP::unc-54 3'UTR; pJRK248, λ DNA]* | This study |
| **BOX786** | *pkc-3(mib78[eGFP-LoxP::AID::pkc-3]) II; mibls49[Pwrt-2::TIR-1::tagBFP2-Lox511::tbb-2-3'UTR, IV:5014740-5014802 (cxTi10882 site))] IV; dlg-1(mib23[dlg-1::mCherry-LoxP]) X; mibEx256[Prps-0::CRY2olig-VH-H(mCherry)::unc-54 3'UTR; Prps-0::CIBN-3xFLAG-MP::unc-54 3'UTR; pJRK248, λ DNA]* | This study |
| **BOX787** | *dlg-1(mib23[dlg-1::mCherry-LoxP]) X; mibEx270[Prps-0::CRY2olig-VH-H(GFP)::unc-54 3'UTR; Prps-0::CIB1-3xFLAG-MP::unc-54 3'UTR; pJRK86(Prps-0::AID::GFP::unc-54 3'UTR), pJRK248, λ DNA]* | This study |
| **BOX788** | *lgl-1(xn103[lgl-1::zf1::mScarlet]) X; mibEx270(Prps-0::CRY2olig-VH-H(GFP)::unc-54 3'UTR; Prps-0::CIB1-3xFLAG-MP::unc-54 3'UTR; pJRK86(Prps-0::AID::GFP::unc-54 3'UTR), pJRK248, λ DNA]* | This study |
| **BOX789** | *erm-1(mib40[erm-1::AID::mCherry]) maph-1.1(mib12[GFP::maph-1.1] l; mibEx256[Prps-0::CRY2olig-VHH(GFP)::unc-54 3'UTR; Prps-0::CIB1-3xFLAG-MP::unc-54 3'UTR; pJRK248, λ DNA]* | This study |
| **BOX790** | *cals[Pges-1::YFP::ACT-5]; mibEx255[Prps-0::mKate2-CRY2olig-VH-H(GFP)::unc-54 3'UTR; Prps-0::CIB1-3xFLAG-MP::unc-54 3'UTR; pJRK248, λ DNA]* | This study |

**Table S2.** Plasmid information.

| **Name** | **Description** | **Function** | **Res. Cassette** | **Reference** |
| --- | --- | --- | --- | --- |
| **pJRK136** | Prps-0::CIBN-3xFLAG::MP::unc-54 3' UTR | CeLINC assay | Ampicillin | This study |
| **pJRK137** | Prps-0::mKate2::CRY2(olig)::VHH(GF-P)::unc-54 3' UTR | CeLINC assay test | Ampicillin | This study |
| **pJRK138** | Prps-0::CRY2(olig)::VHH(GFP)::unc-54 3' UTR | CeLINC assay | Ampicillin | This study |
| **pJRK248** | Prps-0::HygR::unc-54 3'UTR; Psqt-1::sqt-1::sqt-1 3' UTR | co-injection marker | Kanamycin | This study |
| **pJRK249** | Prps-0::CRY2(olig)::VHH(mCherry)::unc-54 3' UTR | CeLINC assay | Ampicillin | This study |
| **pJRK251** | Pwrt-2::CRY2(olig)::VHH(GFP)::unc-54 3' UTR | CeLINC assay | Ampicillin | This study |
| **pJRK252** | Pelt-2::CRY2(olig)::VHH(GFP)::unc-54 3' UTR | CeLINC assay | Ampicillin | This study |
| **pJRK253** | Phsp-16.48::CRY2(olig)::VHH(GF-P)::unc-54 3' UTR | CeLINC assay | Ampicillin | This study |
| **pJRK258** | Prps-0::mKate2::par-6::unc-54 3' UTR | CeLINC assay test | Ampicillin | This study |
| **pJRK259** | Prps-0::mKate2:: par-6(ΔPB1)::unc-54 3' UTR | CeLINC assay test | Ampicillin | This study |
| **pJRK260** | Prps-0::CRY2(olig)::VHH(GFP)::T2A::CIBN-3xFLAG::MP::unc-54 3' UTR | CeLINC assay | Ampicillin | This study |
| **pJRK261** | (empty promoter module with flanking SapI sites)::CRY2(olig)::VHH(GFP)::T2A::CIBN-3xFLAG::MP::unc-54 3' UTR | destination plasmid | Ampicillin | This study |
| **pMLS257** | backbone destination plasmid | destination plasmid | Ampicillin | (Schwartz and Jorgensen, 2016) |
| **pJRK1** | Prps-0 (TGG/ATG overhangs) | donor plasmid | Kanamycin | (Yao et al., 2020) |
| **pJRK151** | Prps-0 (TGG/AAG overhangs) | donor plasmid | Kanamycin | (Yao et al., 2020) |
| **pJRK142** | Phsp-16.48 | donor plasmid | Kanamycin | This study |
| **pJRK143** | Pelt-2 | donor plasmid | Kanamycin | This study |
| **pJRK246** | Pwrt-2 | donor plasmid | Kanamycin | This study |
| **pJRK109** | CRY2(olig) | donor plasmid | Kanamycin | This study |
| **pJRK112** | VHH(GFP) (nanobody) | donor plasmid | Kanamycin | This study |
| **pJRK254** | VHH(mCherry) (nanobody) | donor plasmid | Kanamycin | This study |
| **pJRK107** | CIBN-3xFLAG | donor plasmid | Kanamycin | This study |
| **pJRK108** | MP (multimeric protein) | donor plasmid | Kanamycin | This study |
| **pJRK120** | par-6 cDNA | donor plasmid | Kanamycin | This study |
| **pJRK150** | unc-54 3' UTR (ACG/GTA overhangs) | donor plasmid | Kanamycin | (Yao et al., 2020) |
| **pJRK153** | unc-54 3' UTR (TGC/GTA overhangs) | donor plasmid | Kanamycin | (Yao et al., 2020) |
| **pJRK86** | Prps-0::AID::GFP::unc-54 3' UTR | CeLINC assay test | Ampicillin | This study |
| **pDD375** | mKate2 | donor plasmid | Kanamycin | Gift from Bob Goldstein (Ad-gene plasmid #91825) |

**Table S3.**
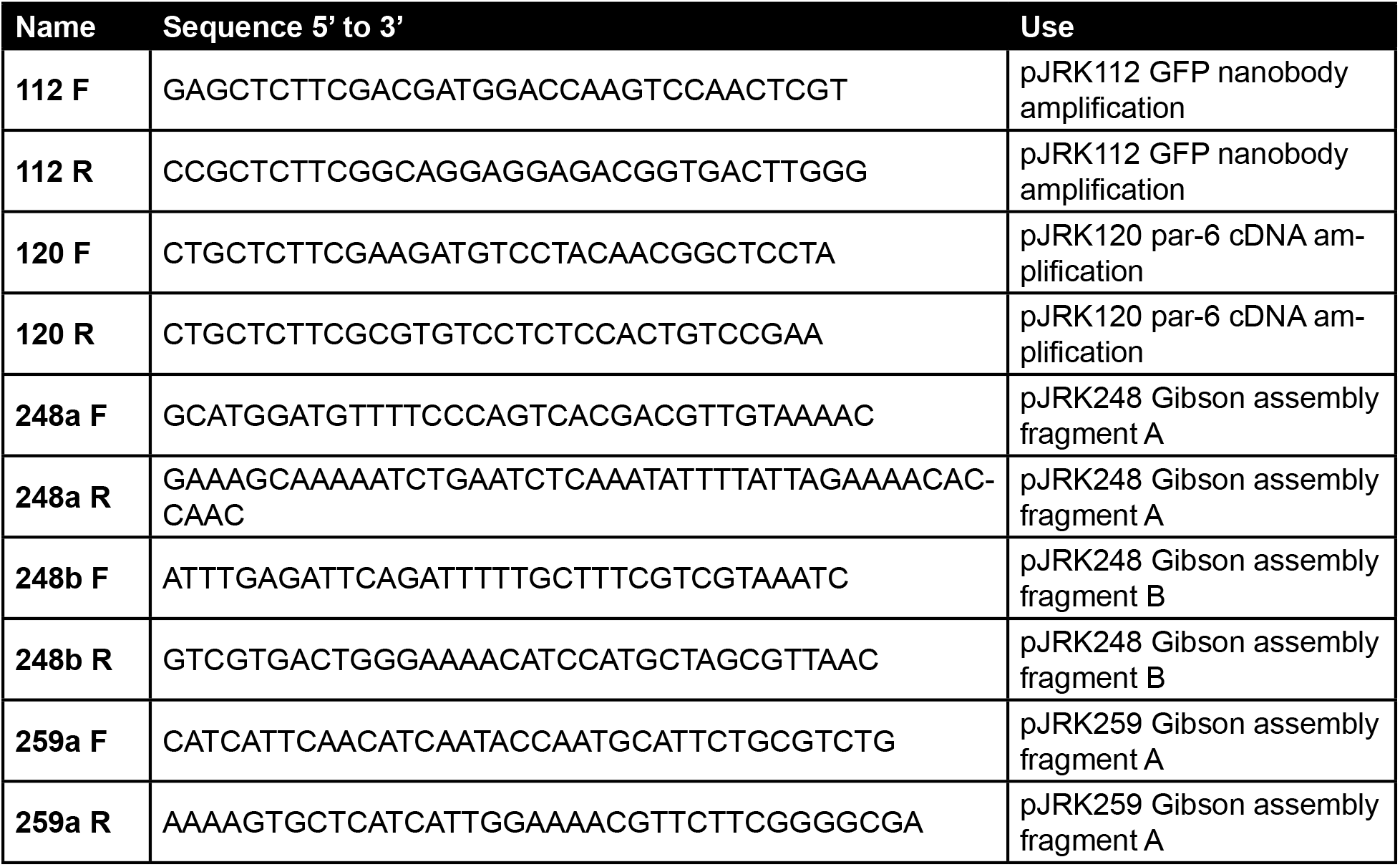
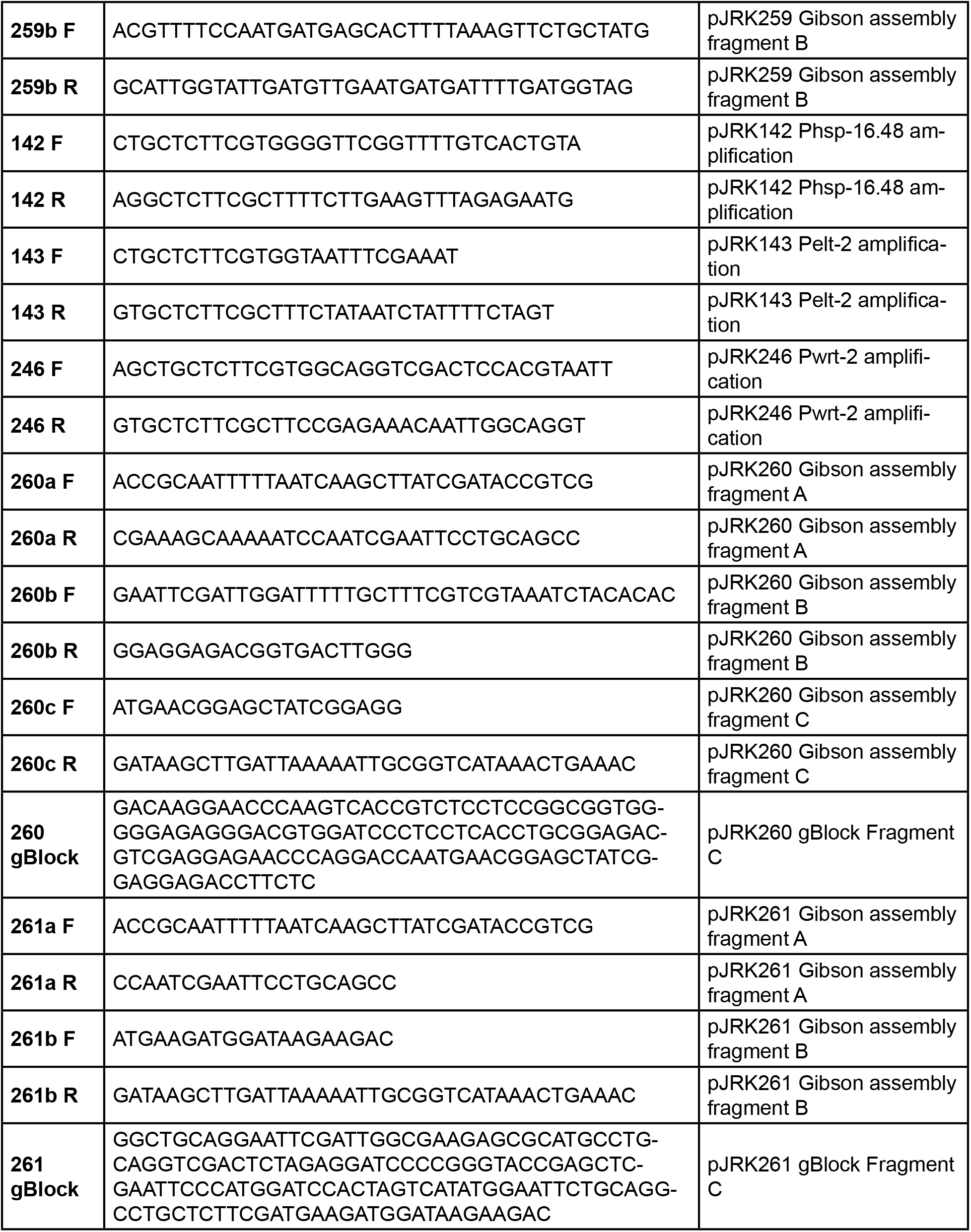
Primers and sequences used for cloning.

**Table S4.**
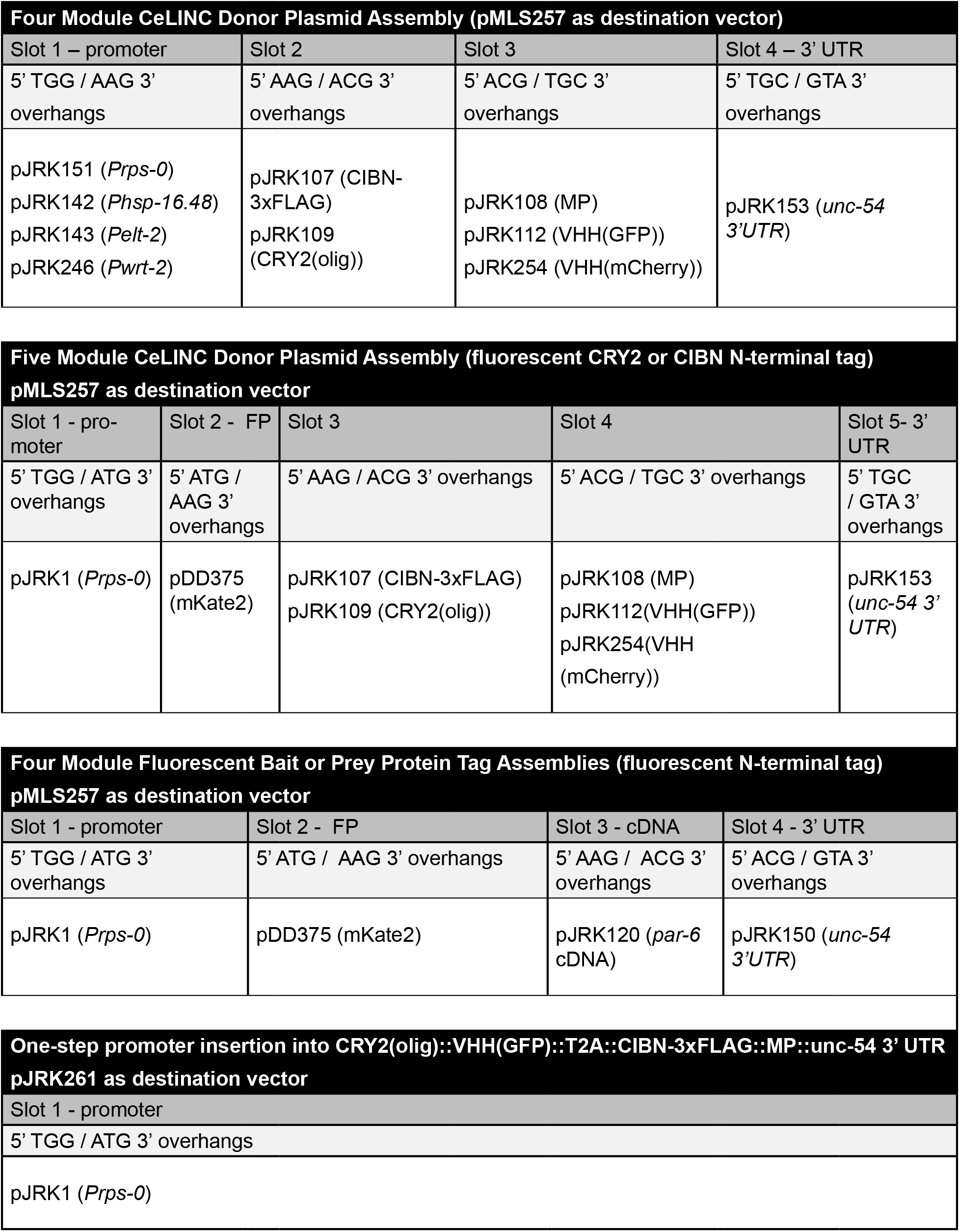
SapTrap assembly information.

| Four Module CeLINC Donor Plasmid Assembly (pMLS257 as destination vector) |  |  |  |
| --- | --- | --- | --- |
| Slot 1 – promoter | Slot 2 | Slot 3 | Slot 4 – 3' UTR |
| 5' TGG / AAG 3' overhangs | 5' AAG / ACG 3' overhangs | 5' ACG / TGC 3' overhangs | 5' TGC / GTA 3' overhangs |
| pJRK151 (*Prps-0*)<br>pJRK142 (*Phsp-16.48*)<br>pJRK143 (*Pelt-2*)<br>pJRK246 (*Pwrt-2*) | pJRK107 (CIBN-3xFLAG)<br>pJRK109 (CRY2(olig)) | pJRK108 (MP)<br>pJRK112 (VHH(GFP))<br>pJRK254 (VHH(mCherry)) | pJRK153 (*unc-54 3'UTR*) |

| Five Module CeLINC Donor Plasmid Assembly (fluorescent CRY2 or CIBN N-terminal tag)<br>pMLS257 as destination vector |  |  |  |  |
| --- | --- | --- | --- | --- |
| Slot 1 - promoter | Slot 2 - FP | Slot 3 | Slot 4 | Slot 5- 3' UTR |
| 5' TGG / ATG 3' overhangs | 5' ATG / AAG 3' overhangs | 5' AAG / ACG 3' overhangs | 5' ACG / TGC 3' overhangs | 5' TGC / GTA 3' overhangs |
| pJRK1 (*Prps-0*) | pDD375 (mKate2) | pJRK107 (CIBN-3xFLAG)<br>pJRK109 (CRY2(olig)) | pJRK108 (MP)<br>pJRK112(VHH(GFP))<br>pJRK254(VHH(mCherry)) | pJRK153 (*unc-54 3' UTR*) |

| Four Module Fluorescent Bait or Prey Protein Tag Assemblies (fluorescent N-terminal tag)<br>pMLS257 as destination vector |  |  |  |
| --- | --- | --- | --- |
| Slot 1 - promoter | Slot 2 - FP | Slot 3 - cDNA | Slot 4 - 3' UTR |
| 5' TGG / ATG 3' overhangs | 5' ATG / AAG 3' overhangs | 5' AAG / ACG 3' overhangs | 5' ACG / GTA 3' overhangs |
| pJRK1 (*Prps-0*) | pDD375 (mKate2) | pJRK120 (*par-6* cDNA) | pJRK150 (*unc-54 3'UTR*) |

| One-step promoter insertion into CRY2(olig)::VHH(GFP)::T2A::CIBN-3xFLAG::MP::unc-54 3' UTR<br>pJRK261 as destination vector |
| --- |
| Slot 1 - promoter |
| 5' TGG / ATG 3' overhangs |
| pJRK1 (*Prps-0*) |

### Supplementary file: Genbank Files

Folder contains annotated GenBank files of the following plasmids:

Supplementary file: Genbank Files Folder contains annotated GenBank files of the following plasmids:
| Donor plasmids |  |
| --- | --- |
| pJRK1 | *Prps-0* |
| pJRK107 | CIBN-3xFLAG |
| pJRK108 | MP |
| pJRK109 | CRY2(olig) |
| pJRK112 | VHH(GFP) |
| pJRK142 | *Phsp-16.48* |
| pJRK143 | *Pelt-2* |
| pJRK150 | *unc-54 3'UTR* |
| pJRK151 | *Prps-0* |
| pJRK153 | *unc-54 3'UTR* |
| pJRK246 | *Pwrt-2* |
| pJRK254 | VHH(mCherry) |

| Other plasmids |  |
| --- | --- |
| pJRK86 | *Prps-0::AID::GFP::unc-54 3' UTR* |
| pJRK136 | *Prps-0::CIBN-3xFLAG::MP::unc-54 3' UTR* |
| pJRK137 | *Prps-0::mKate2::CRY2(olig)::VHH(GFP)::unc-54 3' UTR* |
| pJRK138 | *Prps-0::CRY2(olig)::VHH(GFP)::unc-54 3' UTR* |
| pJRK249 | *Prps-0::CRY2(olig)::VHH(mCherry)::unc-54 3' UTR* |
| pJRK251 | *Pwrt-2::CRY2(olig)::VHH(GFP)::unc-54 3' UTR* |
| pJRK252 | *Pelt-2::CRY2(olig)::VHH(GFP)::unc-54 3' UTR* |
| pJRK253 | *Phsp-16.48::CRY2(olig)::VHH(GFP)::unc-54 3' UTR* |
| pJRK258 | *Prps-0::mKate2::par-6::unc-54 3' UTR* |
| pJRK259 | *Prps-0::mKate2:: par-6(ΔPB1)::unc-54 3' UTR* |
| pJRK260 | *Prps-0::CRY2(olig)::VHH(GFP)::T2A::CIBN-3xFLAG::MP::unc-54 3' UTR* |
| pJRK261 | (empty promoter module with flanking *SapI* sites)::CRY2(olig)::VHH(GFP):<br>:T2A::CIBN-3xFLAG::MP::unc-54 3' UTR |

